# Computational Screening of *Zanthoxylum armatum* DC Phytochemicals for inhibiting Mutant P53 in Throat Cancer Therapy

**DOI:** 10.1101/2024.07.26.605249

**Authors:** Ampi Taring, Senggam Wakhet Singpho, Lobsang Lamu, Bipul Chandra Kalita, Hui Tag, Pallabi Kalita Hui

## Abstract

**Aim:** To investigate the potential of phytochemicals from *Zanthoxylum armatum* DC in targeting mutant p53 as a therapeutic strategy for throat cancer using an in-silico approach.

**Background:** Throat cancer is increasingly associated with mutations in the TP53 gene, leading to dysfunctional p53 proteins. Current synthetic drugs targeting mutant p53 face challenges of toxicity and off-target effects, necessitating the exploration of alternative natural compounds.

**Methods:** We employed molecular docking, molecular dynamics simulations, MMPBSA analysis, and ADMET studies to identify and characterize effective inhibitors of mutant p53 from *Z. armatum* DC phytochemicals.

**Results:** Three compounds – Z3, Z7, and Z33 – demonstrated superior binding affinities to mutant p53 compared to the commercial drug Prima. These phytochemicals induced more compact and stable protein conformations, with Z3 and Z7 notably restricting active site residual motion. MMPBSA analysis confirmed favourable binding energetics, while ADMET studies indicated drug-like properties with potentially lower toxicity than Prima.

**Conclusion:** This study provides compelling evidence for the potential of *Z. armatum* DC phytochemicals, particularly Z3 and Z7, as novel therapeutic agents targeting mutant p53 in throat cancer. These findings offer a foundation for future experimental validation and drug development efforts.

## Introduction

Pharyngeal cancer, colloquially known as throat cancer, is a malignant neoplasm characterized by the aberrant proliferation of cells in the pharyngeal region. This pathological condition arises from somatic mutations that disrupt the delicate balance of cellular homeostasis, leading to uncontrolled growth and potential metastasis. The etiology of pharyngeal cancer is multifactorial, with a complex interplay between genetic predisposition and environmental risk factors. Epidemiological data indicate a concerning global trend in the incidence of pharyngeal cancer. This upward trajectory is attributed to a constellation of modifiable risk factors, including but not limited to, poor nutritional habits, excessive alcohol consumption, tobacco use, and exposure to environmental carcinogens. The synergistic effect of these factors potentiates the risk of carcinogenesis, underscoring the importance of public health initiatives aimed at risk factor mitigation. A comprehensive analysis of global health statistics in 2020 revealed the substantial burden of pharyngeal cancer. Approximately 98,000 new cases were diagnosed worldwide, with a mortality rate of nearly 49%. This translates to an estimated 48,000 deaths annually, highlighting the aggressive nature of this malignancy and the challenges in its management [1]. While pharyngeal cancer accounted for 0.001% of all cancer diagnoses globally, its mortality rate of 0.00051% emphasizes its disproportionate impact on patient survival. The socioeconomic ramifications of pharyngeal cancer are equally alarming. Projections indicate that from 2018 to 2030, the global economic burden attributable to this malignancy will reach a staggering $535 billion USD [2]. This figure encompasses direct healthcare costs, loss of productivity, and the indirect societal impact of premature mortality.

The p53 protein, encoded by the TP53 gene, is a critical regulator of genomic integrity and cellular homeostasis. This multifunctional protein orchestrates a complex network of cellular responses, including cell cycle regulation, DNA repair, and apoptosis induction in response to various stress stimuli [3, 4]. In its wild-type form, p53 functions as a potent tumour suppressor, acting as a guardian against the proliferation of potentially malignant cells [5, 6]. However, in the context of throat cancer, the TP53 gene frequently undergoes mutations, resulting in the production of dysfunctional p53 proteins [7]. These mutant variants not only lose their tumour-suppressive capabilities but can also acquire gain-of-function (GOF) properties that actively promote tumorigenesis. The overexpression of mutant p53 proteins leads to a cascade of cellular dysregulation, including: Disruption of normal cell cycle checkpoints, Impairment of DNA repair mechanisms, Inhibition of apoptotic pathways, Promotion of cellular survival and proliferation, and Enhancement of genomic instability [8–10]. This multifaceted impact of mutant p53 significantly contributes to the initiation, progression, and metastasis of throat cancer. The loss of wild-type p53 function, coupled with the acquisition of oncogenic properties, creates a permissive environment for the accumulation of additional genetic aberrations and the survival of damaged cells. Consequently, targeting mutant p53 has emerged as a promising therapeutic strategy in the management of throat cancer and other p53-mutant malignancies.

Targeting overexpressed mutant p53 protein with synthetic drugs presents a promising therapeutic strategy for treating throat cancer [11, 12]. Compounds such as PRIMA-1, PRIMA-1MET/APR-246 (Eprenetapopt), COTI-2, SAHA, and PEITC have been designed to reactivate the tumour-suppressive functions of p53 or selectively degrade mutant proteins [13, 14]. These drugs aim to inhibit cancer cell proliferation and induce apoptosis, offering a more targeted approach to therapy. However, the application of these synthetic compounds faces significant challenges. A major drawback is the potential for off-target effects, where the drugs might affect normal cells expressing wild-type p53, leading to unintended toxicity and adverse reactions. Clinical trials have reported various side effects associated with these drugs, including nausea, fatigue, peripheral neuropathy, gastrointestinal disturbances, and metabolic changes [5, 15]. Additionally, cancer cells may develop resistance mechanisms over time, limiting the long-term efficacy of these compounds [16]. The limited specificity of some drugs to particular p53 mutations and the complex pharmacokinetics involved in achieving optimal drug concentrations at the tumour site while minimizing systemic exposure further complicate their clinical application [17]. These limitations underscore the need for continued research to optimize drug design, improve delivery methods, and develop combination strategies to enhance efficacy while mitigating side effects in the treatment of throat cancer and other p53-mutant malignancies.

Medicinal plants have emerged as a promising avenue for targeting mutant p53 proteins, offering a complementary and potentially less toxic approach to synthetic drugs in the treatment of throat cancer [18, 19]. This growing interest stems from the diverse array of bioactive compounds found in plants, which can modulate various cellular pathways, including those involving p53. Several phytochemicals have demonstrated significant potential in this regard. Curcumin from turmeric (*Curcuma longa*) has been shown to restore mutant p53 function and induce apoptosis in cancer cells. Its pleiotropic effects on multiple signalling pathways make it a versatile anticancer agent [20]. Epigallocatechin gallate (EGCG) from green tea (*Camellia sinensis*) has dual action, stabilizing wild-type p53 while inhibiting the growth of p53-mutant cancer cells. This unique property makes it particularly interesting for targeting heterogeneous tumour populations [21]. Genistein from soybeans (*Glycine max*) enhances p53 reactivation, promoting cell cycle arrest and apoptosis. Its estrogenic properties may also contribute to its anticancer effects in hormone-responsive cancers [22]. Their multi-target nature may help address the complexity of cancer biology and reduce the likelihood of resistance development. Of particular interest is *Zanthoxylum armatum* DC, a medicinal plant that has shown promising potential in treating throat cancer. Extracts from this plant have demonstrated significant cytotoxic activity against various cancer cell lines, including those related to throat cancer [23, 24]. Methanol extracts and crude saponins from the leaves, bark, and fruits of *Z. armatum* DC have shown the ability to inhibit cancer cell growth and induce apoptosis [25]. These extracts can activate key apoptotic pathways, such as the MAPK pathway, which is crucial for cancer cell death [25]. The multifaceted anticancer properties of *Z. armatum* DC, combined with its potential for lower toxicity, make it a valuable candidate for further investigation in the development of novel throat cancer therapies. This growing body of evidence underscores the potential of phytochemicals, particularly those from *Z. armatum* DC, as a complementary or alternative approach to synthetic drugs in targeting mutant p53 and treating throat cancer.

This study focused on the interaction between mutant P53 and phytochemicals derived from *Zanthoxylum armatum* DC to identify effective inhibitors. Utilizing an in-silico approach, we aimed to discover phytochemicals that could inhibit the overexpression of mutant P53. The methodology involved several computational techniques, starting with ADME-Tox analysis to evaluate the potential of these compounds for clinical application. Further, molecular docking analysis was performed to predict the binding affinity of top hit phytochemicals to the mutant P53 protein. Following this, molecular dynamics simulations were conducted to assess the stability and dynamic behaviour of the protein-ligand complexes. This comprehensive in-silico screening process allowed us to identify phytochemicals with promising inhibitory effects on mutant P53, providing a basis for their potential development as therapeutic agents. The findings suggest that specific phytochemicals from *Zanthoxylum armatum* DC have the potential to inhibit mutant P53 overexpression, which could be significant in developing new treatments for cancers characterized by P53 mutations. This study demonstrates the utility of combining molecular docking, molecular dynamics, and ADMET analysis in identifying and evaluating new potential inhibitors for targeted cancer therapy.

## Methodology

### Collection of plant material and Gas Chromatography-Mass Spectrum Analysis (GC-MS)

*Zanthoxylum armatum* DC fruits were collected from Yazali, Lower Subansiri District of Arunachal Pradesh, India. The fresh fruits of *Zanthoxylum armatum* DC were cleaned thoroughly under tap water and shade dried. The dried fruits were then powdered and soaked in 95% methanol for 12 hrs followed by filtering through Whatman filter No. 41 along with sodium sulphate to remove the sediments and traces of water in the filtrate. The fruit extract was stored and subjected for GC-MS analysis. GC-MS analysis was performed on a GC Clarus 500 Perlin Elmer system comprising an AOC-20i auto sampler and gas chromatograph interfaced to a mass spectrophotometer (GC –MS) instrument using the following conditions: Sample volumes of 1 μl were injected with a splitless mode in to a GC-MS-QP2010 Ultra (Shimadzu) system which consists of TSQ Quantum XLS GC-MS/MS (Thermo Scientific Co.).

### Drug Likeness of top hit phytochemicals

The structures of identified phytochemicals from GC–MS analyses of Zanthoxylum *armatum* DC fruit extract was downloaded from PubChem database (https://pubchem.ncbi.nlm.nih.gov). To evaluate the potential of phytochemicals as lead candidates, we employed Lipinski’s Rule of Five (RO5). This rule provides a set of guidelines that help in the early screening and identification of promising compounds. By calculating the Lipinski values for the phytochemicals under investigation, we could assess their drug-likeness properties. For this purpose, we utilized the SwissADME server (https://www.swissadme.ch/<u>; 28</u>). SwissADME allows the evaluation of the absorption, distribution, metabolism, and excretion (ADME) properties of small molecules, which are crucial factors in determining their suitability as potential drugs. By employing Lipinski’s Rule of Five and the SwissADME server, we could systematically evaluate the phytochemicals under study. This approach enabled us to identify compounds with favourable drug-like characteristics and assess their potential for further development as therapeutic agents, while also comparing them to existing pharmaceuticals already in clinical use.

### ADME-Tox (ADMET) Studies

To assess the potential of the phytochemicals as drug candidates, we performed a comprehensive analysis of their absorption, digestion, metabolism, excretion, and toxicity properties. This evaluation was facilitated by three powerful online tools: SwissADME (https://www.swissadme.ch/; 28) for drug likeness, ProTox 3.0 (https://tox.charite.de/protox3/index.php?site=home; 29) for toxicity analysis. These properties are crucial in determining the pharmacokinetic behaviour of potential drug candidates and their suitability for further development. Using these online resources, we assessed various drug similarity parameters for both the phytochemicals and existing pharmaceuticals. These parameters included lipophilicity (the ability of a compound to dissolve in lipids), solubility (the ability to dissolve in solvents), and drug-likeness score (a measure of a compound’s potential to become a successful drug). By uploading the structural information of the phytochemicals and pharmaceuticals to these servers, we could comprehensively analyse and compare their drug-like properties, toxicity risks, and overall suitability as potential therapeutic agents. We further calculate the boiled-egg plot to evaluate the drug likeness and gastrointestinal absorption of small molecules.

### Molecular Docking

The phytochemicals compounds, which passed the drug-likeness characteristics and ADMET properties were subjected for molecular docking against the binding site of p53 (1tup) protein. The molecular docking process was executed using AutoDock 4.0 [26], a sophisticated algorithm designed for protein-ligand interactions. The receptor’s active site served as the focal point for ligand binding. Our methodology involved preparing the protein structure as the receptor by removing the water molecules, assigning the hydrogen bonds and charges and subsequently minimising the protein [27]. The docking outcomes were meticulously analysed. Given the complex nature of molecular interactions, AutoDock generates an ensemble of potential structures. We rigorously examined the top 10 docked structures, employing a comprehensive selection process that considered not only the numerical scores but also involved visual inspection of the docked configurations. By leveraging this cutting-edge computational tool and employing a meticulous selection process, we aimed to identify the most promising phytochemical candidates for further investigation.

### Molecular Dynamic Simulation

To gain insights into the dynamic behaviour of phytochemicals interacting with p53, we performed Molecular Dynamics (MD) simulations. These simulations were carried out using the GROMACS software package (version 2018.1; 30-32) for a duration of 150 nanoseconds (ns). Before running the simulations, we prepared the protein-ligand complexes by following a specific protocol. CHARMM36 force field was used to define the parameters for the protein and the CGenFF program [33] to generate the topology and parameters for the ligand molecules. The protein-ligand complex was then placed in a cubic water box with dimensions of 5 nm x 5 nm x 5 nm, ensuring sufficient space around the complex to avoid artificial interactions with periodic images. To mimic physiological conditions, we added sodium (Na^+^) and chloride (Cl^-^) ions to the system, neutralizing the overall charge. Energy minimisation of the solvated system was performed to remove any steric clashes or unfavourable contacts, using the steepest descent algorithm with a maximum of 5000 steps or until a convergence threshold was reached. Before the production run, a series of equilibration steps were performed to relax and stabilize the solvent and solute molecules. A 1 ns equilibration run in the NVT ensemble (constant number of particles, volume, and temperature) at 300K, was performed using the Verlet algorithm and the V-rescale thermostat. Following that, a 1 ns equilibration run in the NPT ensemble (constant number of particles, pressure, and temperature) at 1 bar pressure, was carried out using the Berendsen pressure coupling algorithm. After equilibration, the production MD simulations for 150 ns was performed, using the leap-frog algorithm with a 2-femtosecond time step for integrating the equations of motion. The van der Waals interaction cutoff was set to 1.2 nm with a smooth switching function and a 1.2 nm cutoff for Coulombic interactions using the Particle Mesh Ewald (PME) method. Constant temperature and pressure were maintained using the Nosé-Hoover thermostat and the Parrinello-Rahman barostat, respectively. The system coordinates were stored every 20 picoseconds during the 150 ns production run for further analysis [27, 34]. Upon completing the simulations, the trajectories were analysed to understand the structural implications of the protein-ligand interactions

### Molecular Mechanics – Poisson-Boltzmann Surface Area (MMPBSA)

Calculating the binding free energy (Δ*G*_*bind*_) between a protein and its ligand is a crucial step in understanding the molecular interactions that govern protein-ligand complexes. In our study, we employed the widely accepted Molecular Mechanics/Poisson-Boltzmann Surface Area (MM/PBSA) methodology to estimate the binding free energy of these complexes. We utilized the fastDRH webserver [35] for this purpose. The binding free energy (Δ*G*_*bind*_) in the MM/PBSA method is computed as the difference between the free energy of the complex (Δ*G*_*complex*_) and the sum of the free energies of the isolated protein (Δ*G*_*protein*_) and ligand (Δ*G*_*ligand*_). Mathematically, it can be expressed as:

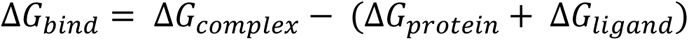

In this equation, Δ*G*_*complex*_ represents the free energy of the protein-ligand complex, Δ*G*_*protein*_corresponds to the free energy of the protein in its unbound state, and Δ*G*_*ligand*_ denotes the free energy of the ligand in its unbound state. By calculating the binding free energy, we can gain insights into the favourability and strength of the interactions between the protein and its ligand. A more negative value of Δ*G*_*bind*_indicates a stronger binding affinity between the two molecules, suggesting a more stable and energetically favourable complex formation.

## Results

The GC-MS analysis of Zanthoxylum *armatum* DC fruits revealed the presence of 116 compounds (phytochemical constituents) and out of which 43 compounds were shortlisted for molecular docking based on organic properties. The drug-likeness property of a compound characterizes its potential to act as a drug. To evaluate this property for the top hit phytochemicals, we employed two powerful online tools: the SwissADME server. These servers enabled us to calculate the drug-likeness and toxicity properties of the selected compounds. Among the top hits, all the ligands followed Lipinski’s rule, indicating favourable drug-like properties, as they showed no violations of the rule’s parameters. We calculated the Lipinski’s factor for the 43 phytochemicals of *Zanthoxylum armatum* DC (Supplementary Table 1) and selected the phytochemicals following all the Lipinski’s rule. After performing the initial screening, we further performed an LD50 screening analysis, using ProTox 3.0 server (https://tox.charite.de/protox3/index.php?site=home; 29), to select the non-toxic phytochemicals. LD50 is a toxicological property that measures the lethal dose of a given substance. LD50 corresponds to the dose of a substance required to kill half the members of a tested population. We selected the phytochemicals showing an LD50 of more than or equal to 2000 mg/kg. Table 1 corresponds to the top 18 phytochemicals left after performing the initial screening (Figure 1). The details about other phytochemicals are provide in Supplementary Table1.

**Figure 1:**
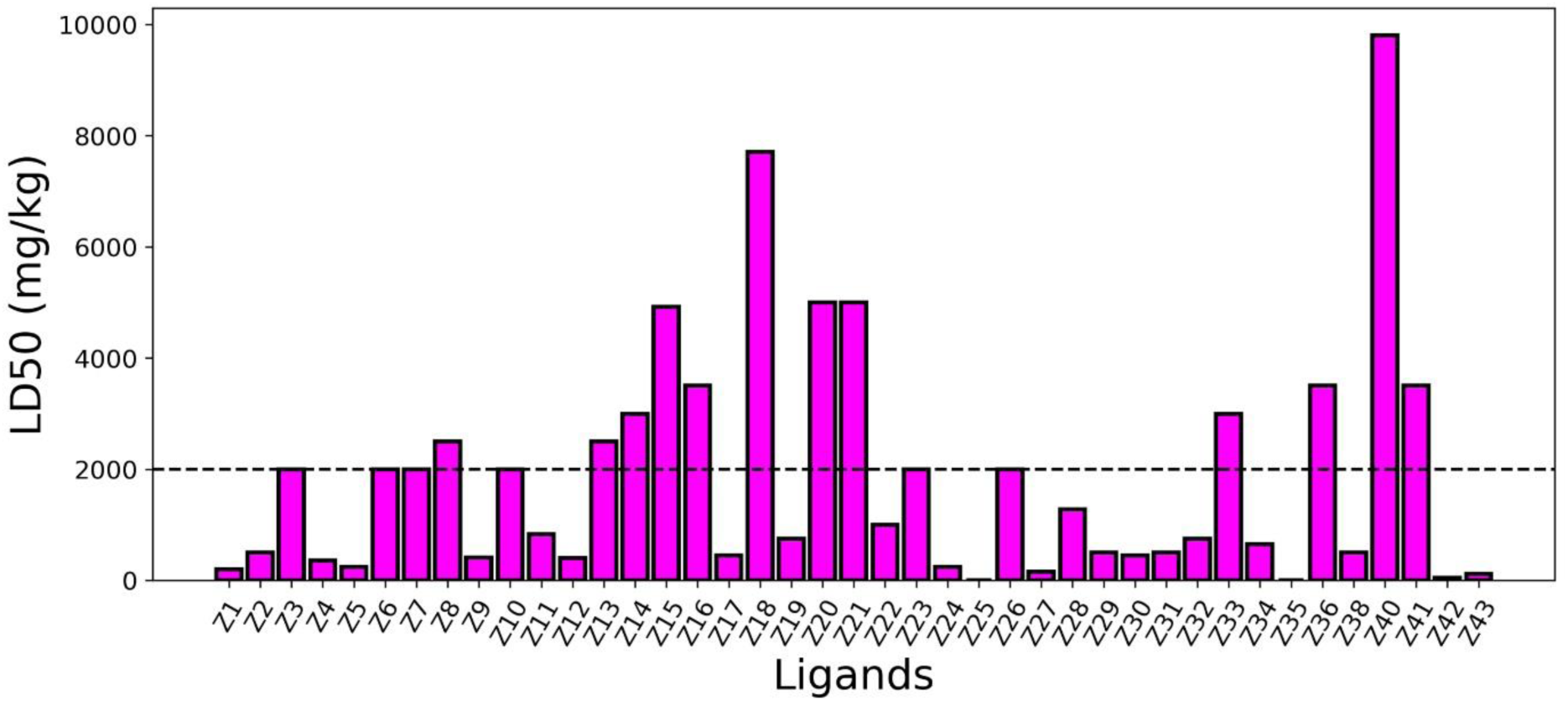
LD50 Screening of the phytochemicals of *Zanthoxylum armatum.* The horizontal dashed line represents the screening cutoff (≥ 2000 mg/kg).

**Table 1:** Details of Phytochemicals selected after Initial Screening.

| Drugs | Lipinski's Rule of Five |  |  |  | LD50 (mg/kg) |
| --- | --- | --- | --- | --- | --- |
| | Mass ($\leq 500$ Da) | HBD ($\leq 5$) | HBA ($\leq 10$) | LogP ($< 5$) | |
| Prima | 185.22 | 2 | 4 | -0.16 | 1500 |
| Z3 | 286.41 | 2 | 2 | 3.99 | 2000 |
| Z6 | 273.42 | 1 | 1 | 4.3397 | 2000 |
| Z7 | 394.59 | 4 | 4 | 3.35 | 2000 |
| Z8 | 187.19 | 2 | 1 | 2.2657 | 2500 |
| Z10 | 329.396 | 2 | 5 | 2.8475 | 2000 |
| Z13 | 322.36 | 2 | 4 | 4.3799 | 2500 |
| Z14 | 276.419 | 1 | 2 | 3.5690 | 3000 |
| Z15 | 186.209 | 2 | 2 | 2.7648 | 4920 |
| Z16 | 261.408 | 1 | 2 | 3.4648 | 3500 |
| Z18 | 251.282 | 2 | 4 | 1.0025 | 7710 |
| Z20 | 166.219 | 0 | 2 | 1.6991 | 5000 |
| Z21 | 372.502 | 3 | 6 | 1.7901 | 5000 |
| Z23 | 296.414 | 2 | 2 | 3.9993 | 2000 |
| Z26 | 329.396 | 2 | 5 | 2.8475 | 2000 |
| Z33 | 275.347 | 0 | 4 | 1.1897 | 3000 |
| Z36 | 261.408 | 1 | 2 | 3.4648 | 3500 |
| Z40 | 192.167 | 5 | 5 | -2.3213 | 9800 |
| Z41 | 171.195 | 3 | 4 | -1.6753 | 3500 |

*HBD – Number of Hydrogen Bond Donor*
*HBA – Number of Hydrogen Bond Acceptor*

When assessing a compound’s potential as a drug candidate, lipophilicity (LogP) is a crucial factor to consider. The LogP value influences the rate at which drug molecules are absorbed by the body. Generally, drugs with higher LogP values tend to be more hydrophobic and non-polar, which is often preferred for intracellular medications. However, the most desirable properties of a drug may vary depending on various factors. The LogP values for the top hit phytochemicals depicted a good range of values (Table 1). Another important parameter is the Synthetic Accessibility (SA) score, which indicates how easy it is to synthesize a compound. An SA value of 1 suggests that the compound can be readily synthesized, while a value of 10 indicates a difficult synthesis process. The phytochemicals under investigation had SA values ranging from 3.60 to 5.20, indicating a moderate synthesis process similar to Prima (Table 2). Especially Z3 showed a similar SA score as that of Prima. The Bioavailability Score (BS) represents the amount of drug that will reach the desired target site after administration into the body. All the top hit ligands showed a similar bioavailability score of 0.55 (Table 2). Even though the SA score of Z33 was higher than Prima, it acquired a better position in the boiled egg plot representing a Blood Brain Barrier permeation (falling in yellow region) and Pgp effluent (blue colour) (Figure 2).

**Figure 2:**
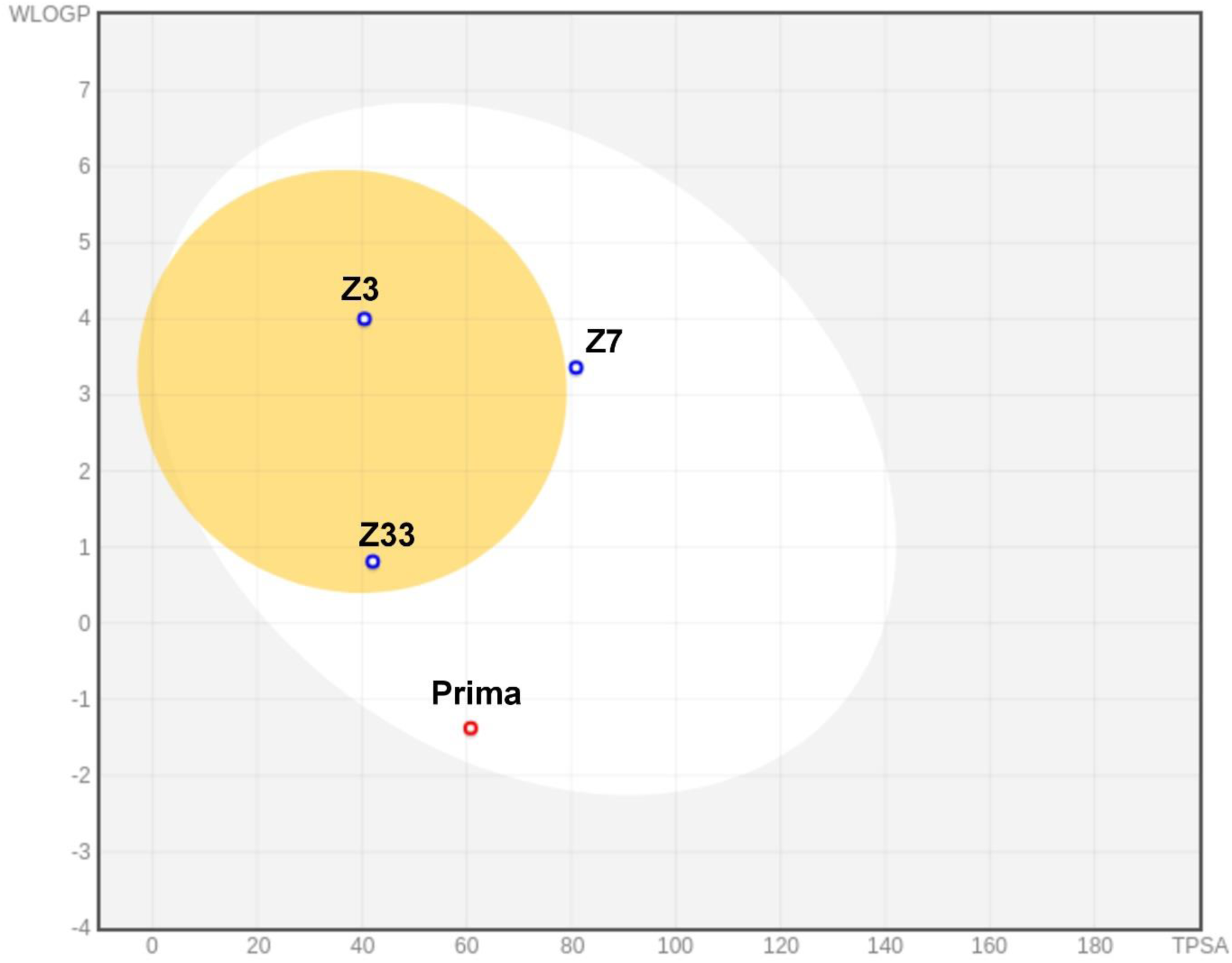
Boiled Egg plot for the top hit phytochemicals and commercial drug.

**Table 2:** ADMET analysis of selected ligands.

| Property | Prima | Z3 | Z7 | Z33 |
| --- | --- | --- | --- | --- |
| GI Absorption | High | High | High | High |
| Bioavailability Score | 0.55 | 0.55 | 0.55 | 0.55 |
| Synthetic Accessibility | 3.08 | 3.60 | 5.20 | 5.09 |

### Toxicity

Toxicity analysis was carried out to understand the toxic characteristics of the top hit ligands by utilising the ProTox 3.0 [29]. We found that the commercial drug Prima turned out be positive for both neurotoxicity and respiratory toxicity (Figure 3a). Among the top ligands, all the three ligands were toxic against respiratory system. Further Z7 and Z33 turned out to be toxic against cardiotoxicity and neurotoxicity respectively. In the case of toxicity end points (Figure 3b), Prima showed positive toxicity against Blood Brain Barrier and Clinical toxicity. Among the top hit ligands, Z33 showed a higher toxicity against different end points such as Carcinogenicity, BBB, Ecotoxicity and Clinical toxicity. Z3 and Z7 showed positive toxicity against Immunotoxicity, Ecotoxicity, Clinical toxicity and Immunotoxicity, BBB and Clinical toxicity respectively. The toxicity behaviour of Z7 resembled closely with that of Prima. In the case of Tox21 Stress pathway (Figure 3c), except for Z3 all the phytochemicals and Prima showed no toxic behaviour. Whereas Z3 showed a positive toxicity against p53 and Mitochondrial Membrane Potential (MMP). In the case of Tox21 Nuclear Receptor signalling pathway (Figure 3d), except for Z3 all the phytochemicals and Prima were nontoxic for each of the receptors. Z3 showed a positive toxicity against Androgen Receptor (AR), Androgen Receptor Ligand Binding Domain (ARLBD), Estrogen Receptor Alpha (ER), and Estrogen Receptor Ligand Binding Domain (ERLBD). All of the phytochemicals and Prima showed toxicity against molecular initiating events (Figure 3e). Furthermore, except for Z7, both the phytochemicals and Prima, showed toxicity against CYP2C9 (showed by Z3), CYP2D6 (showed by Z33) and CYP2C19 and CYP2C9 (showed by Prima) respectively (Figure 3f). This analysis revealed a similar and potential less toxic behaviour of Z7 as a drug candidate. But an important thing that has to be considered is that the results are obtained from models trained on experimental data and the results are probabilistic. One needs to perform experimental examination to further confirm the actual toxicity. Nevertheless, this analysis establishes a structured pipeline to select the less non-toxic compounds for further analysis.

**Figure 3:**
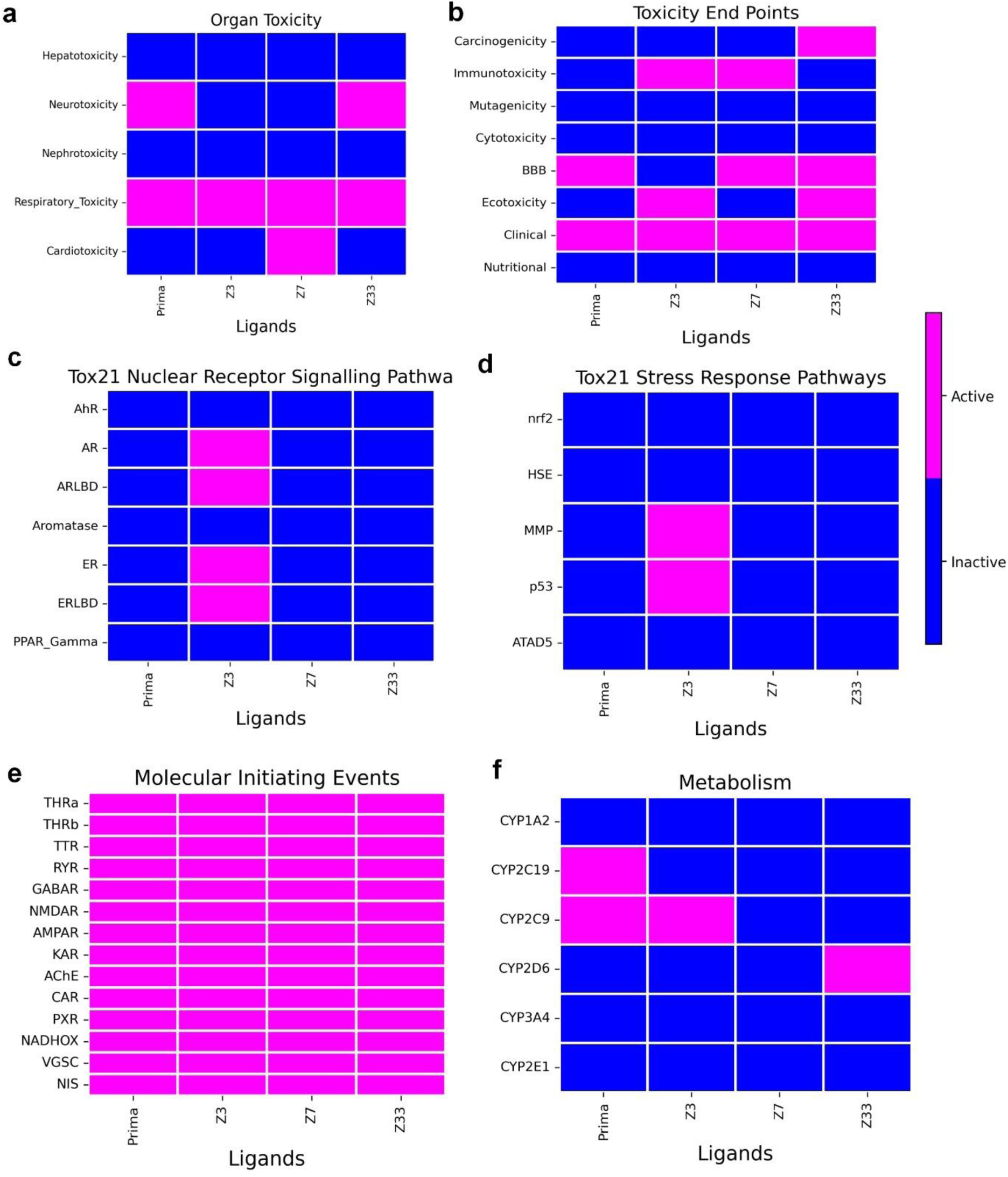
Toxicity analysis of Prima and top hit phytochemicals.

### Molecular Docking

Our investigation employed AutoDock 4.0 [26] to conduct comprehensive molecular docking studies between the ligands and the receptor. The receptor’s active site served as the crucial binding interface for the ligands. To quantify the binding affinity, we utilized the binding energy as a parameter, following established protocols in the field [36, 37]. Our rigorous screening process identified three phytochemicals (Figure 4) that demonstrated significant binding to the protein’s active site. All the top hit phytochemicals showed a better binding energy with the protein than Prima (Table 3).

**Figure 4:**
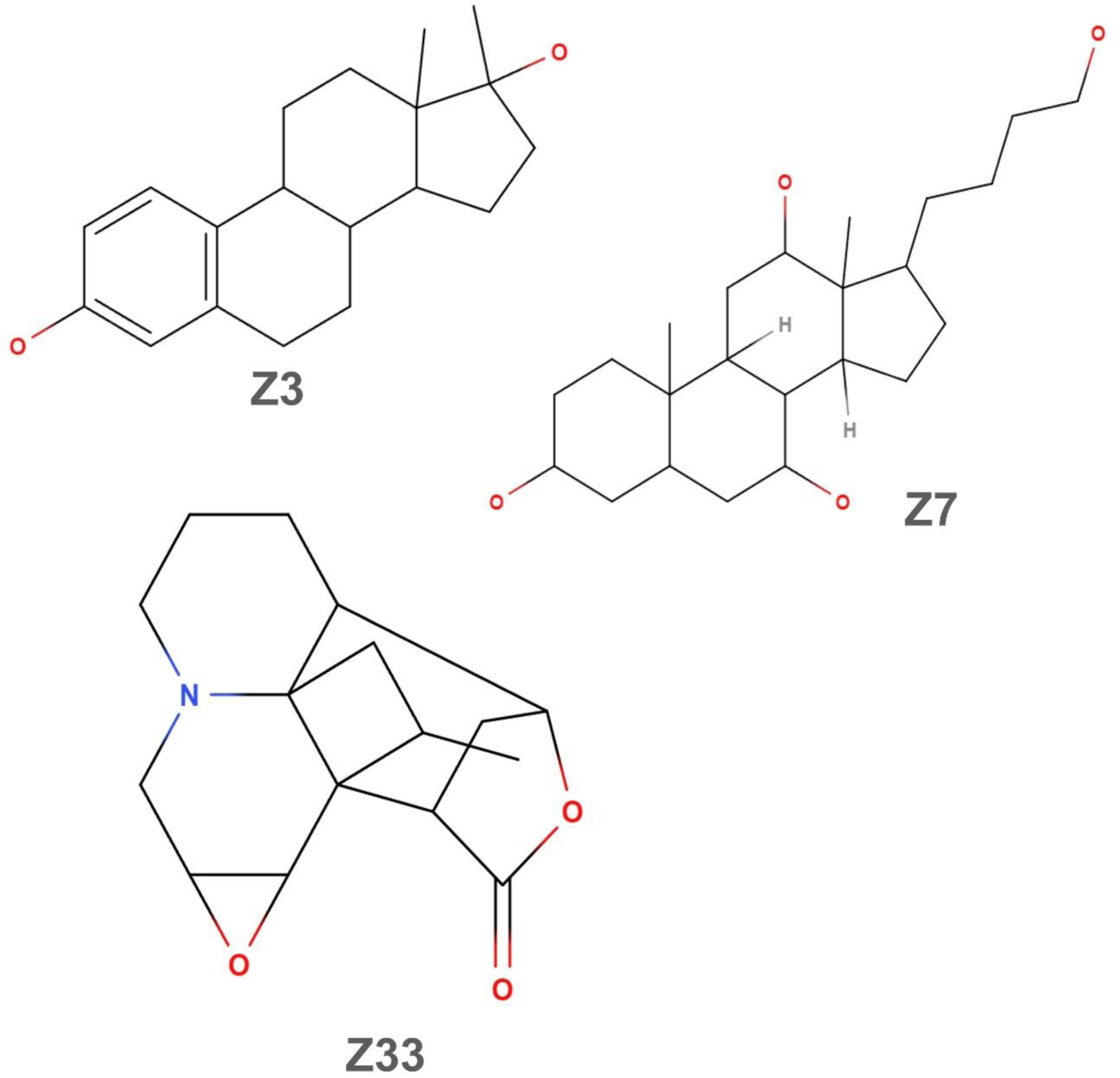
Top hit phytochemicals of *Zanthoxylum armatum* DC selected after docking.

**Table 3:**
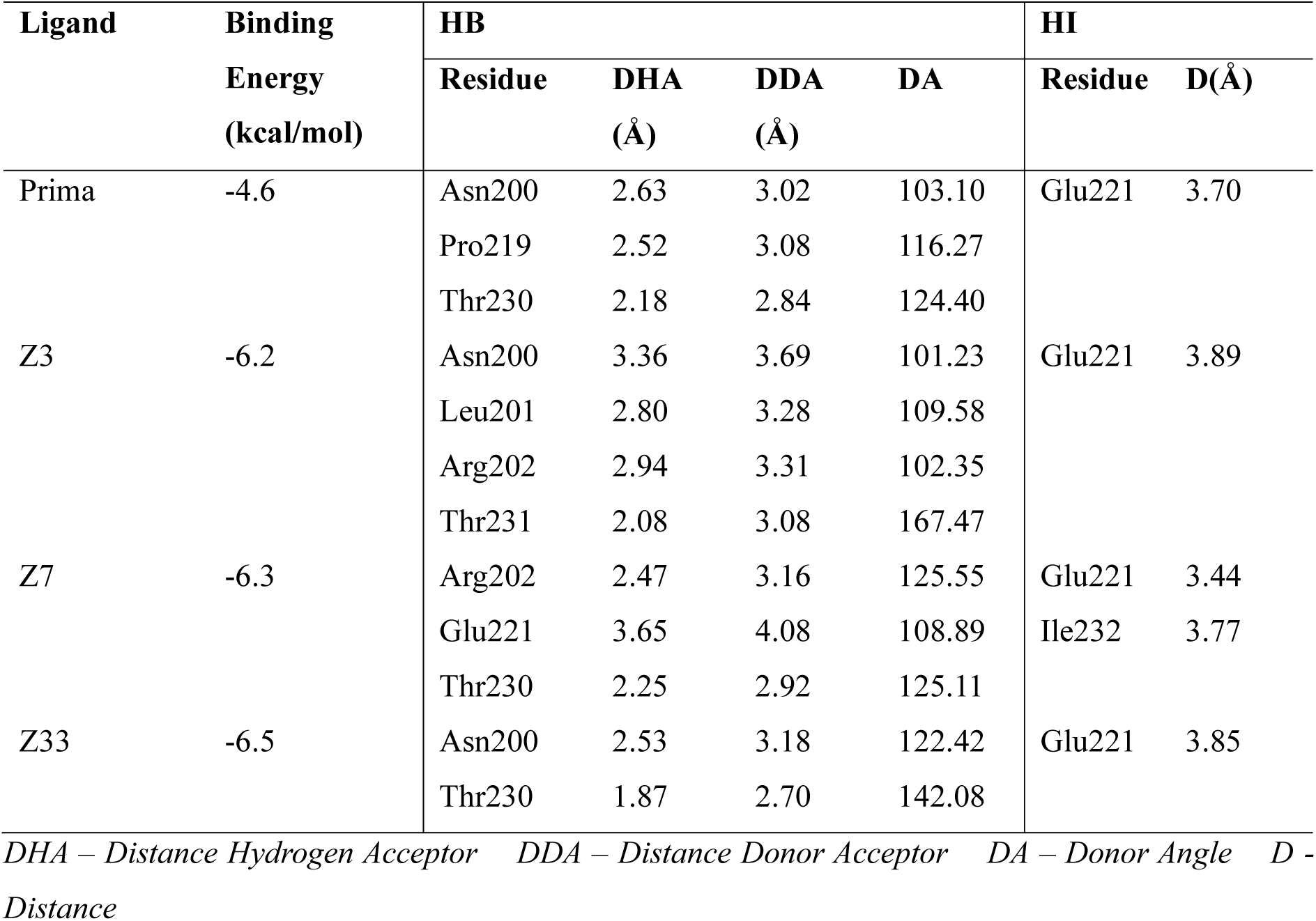
Molecular Docking Analysis of Prima and Top hit phytochemicals.

Prima showed hydrogen bonding with three residues: Asn200, Pro219 and Thr230 with a binding energy of –4.6 kcal/mol. Further it showed a hydrophobic interaction with Glu221. Albeit being a commercial drug [38], Prima shows a poor binding affinity with the protein compared to the top hit phytochemicals. Z3, our first lead compound, formed an intricate network of hydrogen bonds with four residues involving two key residues: Asn200, Leu201, Arg202, Thr231 (Table 3). This complex interaction yielded a remarkable binding energy of – 6.2 kcal/mol (Figure 5(b)). The hydrogen bond distances ranged from 2.08 to 3.36 Å, with notably strong interactions observed with Thr231 (2.08 Å). These short-range interactions likely contribute significantly to the stability of the protein-ligand complex. Z7, our second lead compound, demonstrated an impressive binding characteristic. It formed hydrogen bonds with three residues: Arg202, Glu221 and Thr230 (Figure 5(c)). This extensive network of interactions resulted in a binding energy of −6.3 kcal/mol. Particularly strong hydrogen bonds were observed with Arg202 (2.47 Å) and Thr230 (2.25 Å). Additionally, Z7 engaged in hydrophobic interactions with Glu221 and Ile232 enhancing its binding stability. Z33, our third lead compound, exhibited a unique binding profile. It formed hydrogen bonds with two residues: Asn200 and Thr230 (Figure 5(d)). This interaction pattern yielded a binding energy of −6.5 kcal/mol. Notably, Z33 formed exceptionally strong hydrogen bonds with Thr230 (1.87 Å). The compound also engaged in hydrophobic interactions with Glu221, contributing to its overall binding affinity. This comprehensive analysis identified Z3, Z7 and Z33 as top-performing phytochemicals, exhibiting superior binding affinities for the active pocket of mutant p53. These findings highlight the potential of these compounds as lead molecules.

**Figure 5:**
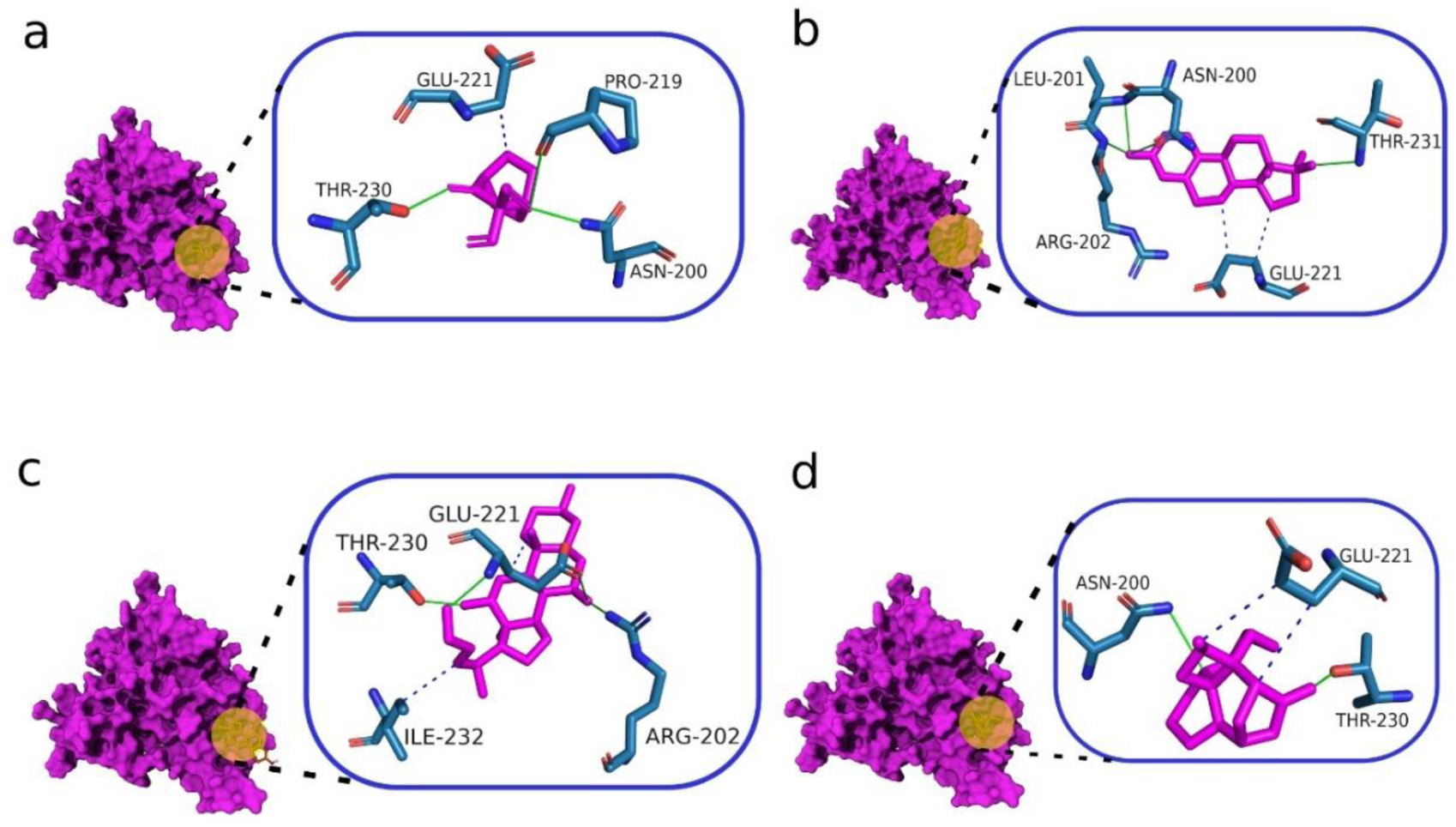
Molecular Docking Interaction of mutant p53 (magenta colour protein; surface view) with (a) Prima (b) Z3 (c) Z7 and (d) Z33. The yellow circle highlights the binding region of respective ligands in the surface view. The zoomed in image shows the residual interaction. Hydrophobic Interaction (green line) and Hydrophobic Interaction (blue dashed line)

### Ligand and Receptor Preparations

To gain insights into the dynamic behaviour of the p53 protein-ligand complexes, we performed a 150 nanosecond (ns) molecular dynamics (MD) simulation. During the simulations, we analysed relevant properties such as root mean square deviation (RMSD), root mean square fluctuation (RMSF), hydrogen bonding (H-bond), solvent accessible surface area (SASA), and radius of gyration. These analyses allowed us to explore the dynamics of the phytochemical-protein complexes. By performing extensive MD simulations and analysing various properties, we gained valuable insights into the dynamic behaviour of the protein-ligand complexes, the stability of the binding modes, and the potential impact of the phytochemicals on the protein’s structure.

### Root Mean Square Deviation

The Root Mean Square Deviation (RMSD) is a valuable metric for assessing the stability of protein-ligand complexes during Molecular Dynamics (MD) simulations. It measures the average deviation of the atomic positions from a reference structure over the course of the simulation trajectory. In our study, we analysed the RMSD profiles of the top hit ligands in complex with the protein to evaluate their binding stability throughout the 150 ns MD simulation. The results demonstrated that most of the top hit ligands exhibited stable binding interactions with the protein, as evidenced by their RMSD fluctuations within acceptable limits and following a similar RMSD fluctuation pattern as that of protein (Figure 6, black). Specifically, the RMSD fluctuations for Z3 and Z33 were within the permissible ranges of 0.1–0.21 nm and 0.1–0.25 nm, respectively (Figure 6, green/magenta). Prima, on the other hand, displayed slightly higher fluctuations after 80 ns, ranging from 0.15 to 0.25 nm (Figure 6, red), suggesting a moderate degree of conformational flexibility during the simulation. Interestingly, Z7 exhibited a distinct behaviour. Initially, it demonstrated a stable fluctuation until 120 ns. However, beyond this point, the RMSD increased significantly during the 120–130 ns timeframe, indicating a conformational change in the protein-ligand complex. Subsequently, the RMSD returned back to common fluctuation range, reaching an average value of 0.18 nm after 135 ns until the end of the simulation (Figure 6, blue). This observation suggests that Z7 may have undergone a conformational rearrangement or potential dissociation from the binding pocket during the later stages of the simulation. In contrast, Z3 exhibited a lower fluctuation pattern over the entire MD simulation. This behaviour indicates that the protein-ligand complex attained a stable conformation throughout the simulation. The observed RMSD profiles suggest that the binding of the respective ligands induces distinct conformational changes in the protein throughout the MD simulation. These conformational rearrangements may be crucial for achieving optimal binding interactions and stabilizing the protein-ligand complexes.

**Figure 6:**
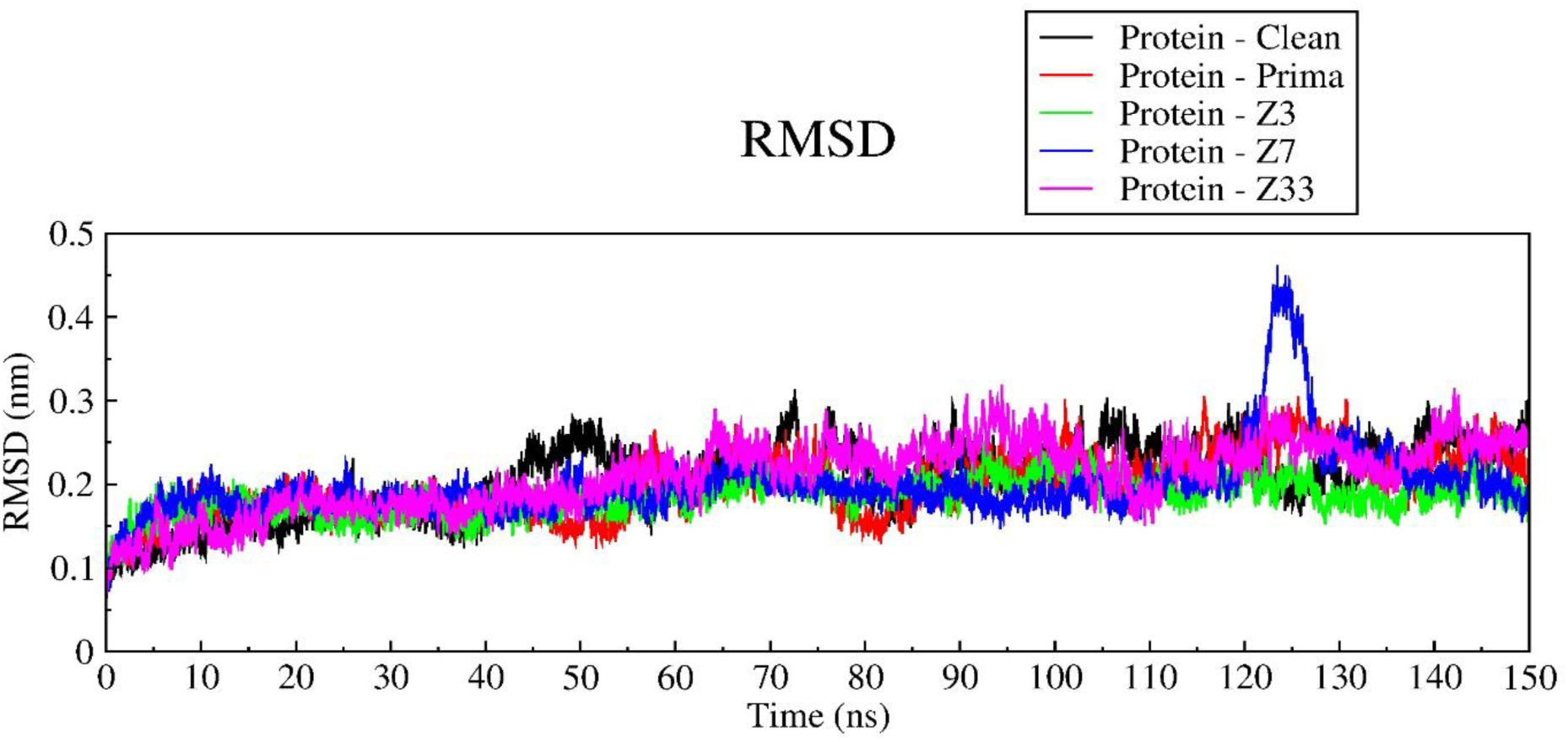
Root Mean Square Fluctuation of Protein without any ligand (Black), Prima (red) and top hit phytochemicals: Z3 (Green), Z7 (Blue) and Z33 (Magenta) for a Molecular Dynamic Simulation of 150 ns.

### Radius of Gyration

The Radius of Gyration (RoG) is a crucial parameter that provides insights into the stability, compactness, and folding state of proteins (Sangeet et al., 2022). In our study, we analysed the RoG profiles of the protein in complex with the top hit ligands to evaluate their impact on the protein’s structural integrity during the 150 ns MD simulation. The results demonstrated that upon binding of the top hit ligands, the protein remained relatively compact and stable throughout the simulation trajectory (Figure 7). Notably, the binding of Prima induced RoG fluctuations in the range of 1.62-1.67 nm (Figure 7, red). Interestingly, the fluctuation marginally decreased after 120 ns, indicating an increase in the protein’s compactness during the latter stages of the simulation. In contrast, the top hit ligands exhibited a more favourable gyration behaviour compared to Prima. Z3, Z7, and Z33 (Figure 7, green: blue: magenta) displayed similar RoG profiles, with fluctuations ranging from 1.62 to 1.69 nm. This observation suggests that these ligands maintained the protein’s structural integrity and compactness throughout the simulation as they resembled the RoG profile of protein (Figure 7, black). Notably, Z33 exhibited a higher RoG value compared to the other top hit ligands, indicating a more extended conformation of the protein-ligand complex (Figure 7, magenta). Strikingly, after 85 ns, the RoG fluctuation of the protein increased significantly, suggesting that the binding of Z33 caused a loss of structural integrity in the protein during the later stages of the simulation. Eventually post 120 ns, the RoG value decreased and then increased again suggesting that the binding of Z33 causes higher structural disintegration in the protein. Z3, on the other hand, showed a good resemblance in RoG as that of Prima suggesting that the ligand causes the protein to remain intact throughout the simulation (Figure 7, green). The observed RoG profiles provide valuable insights into the impact of ligand binding on the protein’s structural stability and compactness. The favourable RoG behaviour observed for most top hit ligands supports their potential as viable candidates for further development, as they maintained the protein’s structural integrity throughout the simulation. However, the distinct behaviour exhibited by Z3, warrants further investigation to understand the underlying dynamics and their implications for binding affinity and stability.

**Figure 7:**
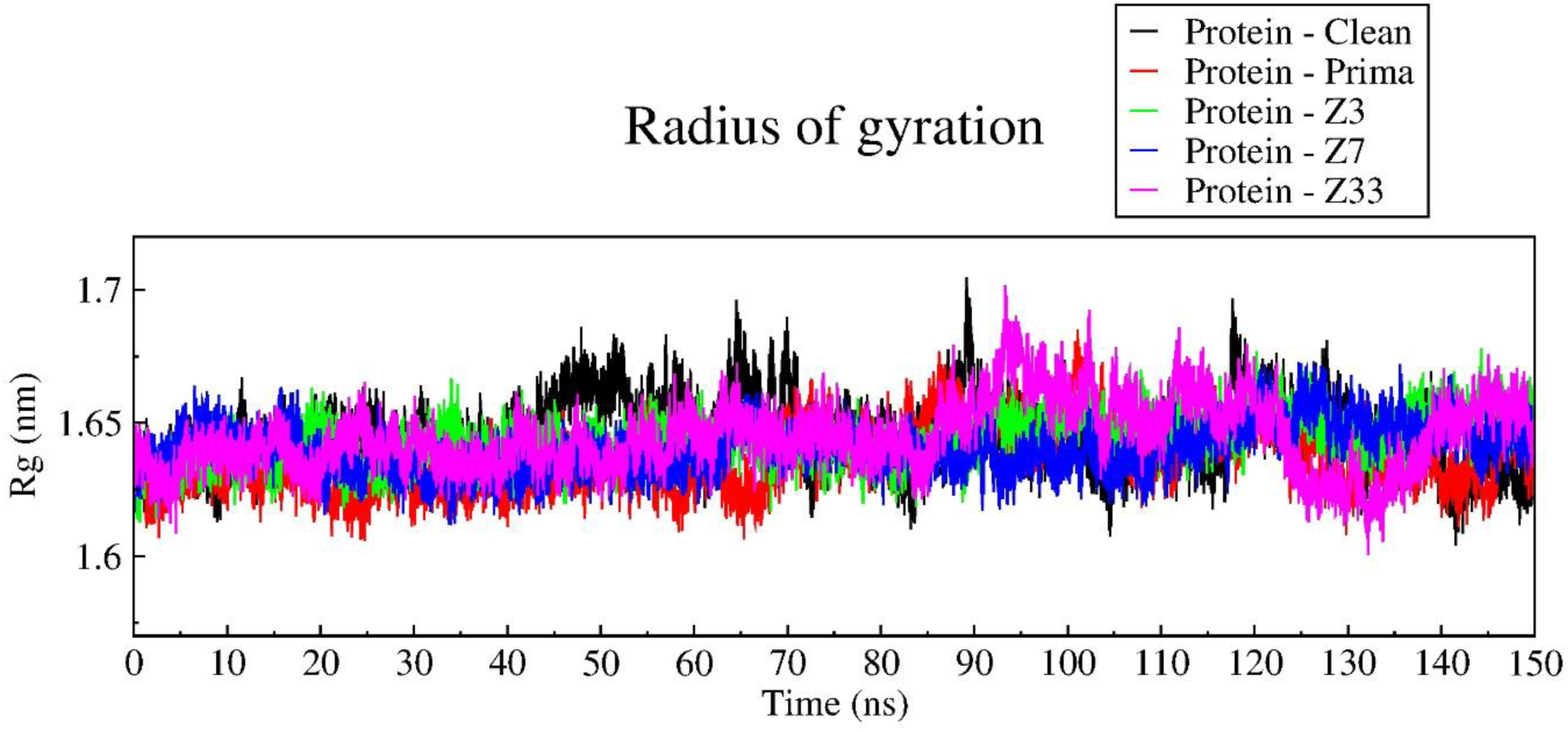
Radius of Gyration of Protein without any ligand (Black), Prima (red) and top hit phytochemicals: Z3 (Green), Z7 (Blue) and Z33 (Magenta) for a Molecular Dynamic Simulation of 150 ns.

### Solvent Accessible Surface Area

The Solvent Accessible Surface Area (SASA) analysis provides valuable insights into the surface area of a protein that is exposed to solvent molecules. A higher SASA value indicates a relative expansion of the protein structure, while a lower value suggests a more compact conformation. In this study, we analysed the SASA profiles of the mutant p53 receptor in complex with various ligands to understand the impact of ligand binding on the protein’s conformational dynamics and solvent exposure. As illustrated in Figure 8, when mutant p53 interacts with the reference ligand Prima (red curve), the SASA value remains stable till 50 ns after which the SASA increases and reaches a maximum value of 122.5 nm^2^ at 100 ns and remained at a higher value throughout the 150 ns simulation, ranging from 113 to 126 nm^2^. Interestingly, all the top hit ligands, except for Z33, induced a more compact conformation of mutant p53 compared to Prima, as evident from their lower SASA values. Among the top hit phytochemicals, Z3 (Figure 8, green) and Z7 (Figure 8, blue) exhibited a constant SASA value in the range of 102 to 115 nm^2^, suggesting that these ligands maintained a highly compact protein conformation throughout the simulation. On the other hand, Z33 (Figure 8, magenta) displayed a constant SASA profile until 50 ns, after which the SASA values started to increase and fluctuate slightly at a higher value than Z3 and Z7. This observation indicates that the binding of Z33 initially promoted a compact conformation, but later induced a slight expansion of the protein structure. The SASA profile demonstrates that the binding of Prima causes the mutant p53 protein to adopt an unstructured conformation, as evidenced by the higher SASA values. In contrast, the top hit ligands are capable of inducing a more compact protein structure, as reflected by their lower SASA values. These findings suggest that the top hit phytochemicals, particularly Z3 and Z7, have the potential to stabilize the p53 receptor in a more compact and structured conformation, which may have implications for its function and interactions with other biomolecules. The differential effects of the ligands on the protein’s conformational dynamics highlight the importance of considering SASA analysis in the evaluation of potential drug candidates and their impact on target protein structures.

**Figure 8:**
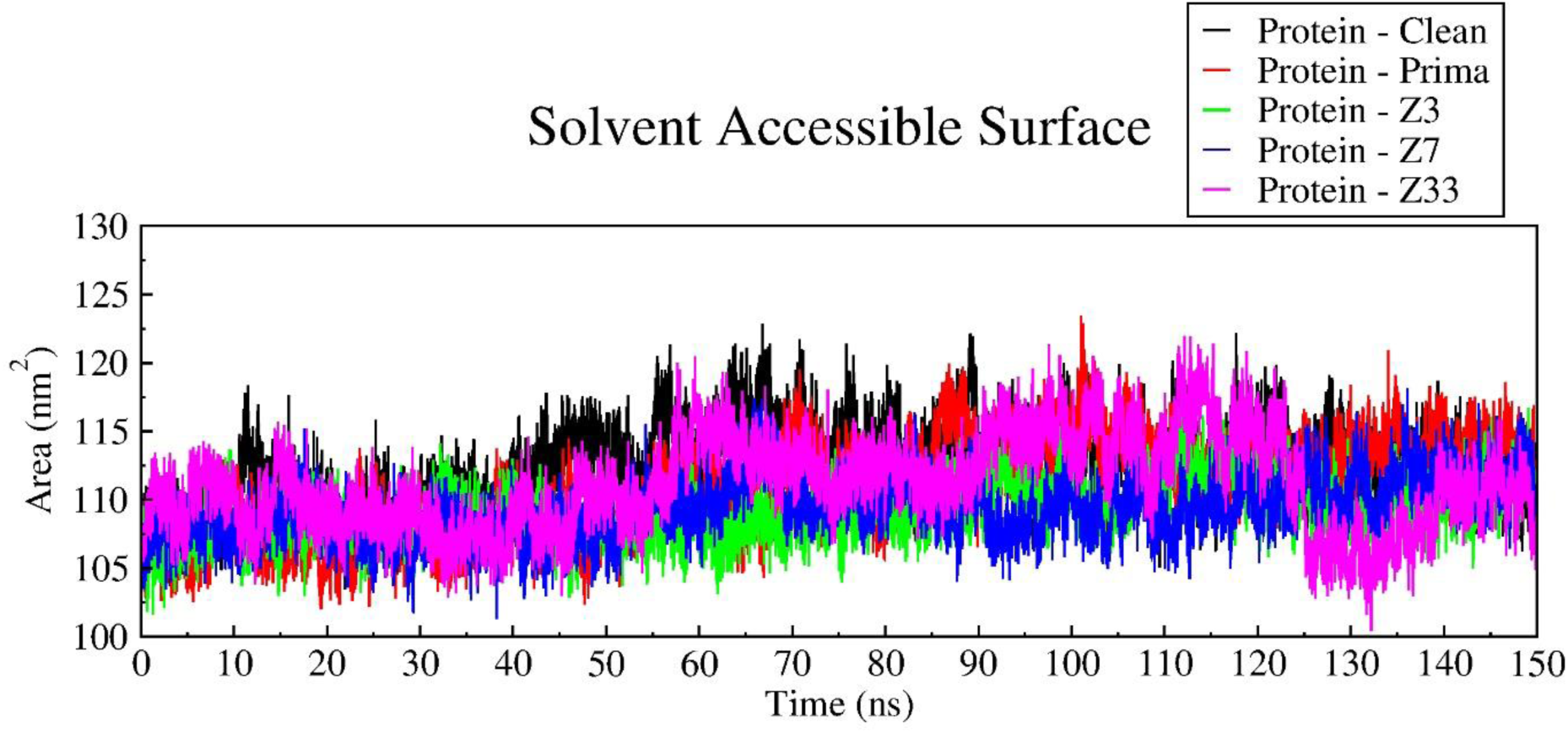
Solvent Accessible Surface Area of Protein without any ligand (Black), Prima (red) and top hit phytochemicals: Z3 (Green), Z7 (Blue) and Z33 (Magenta) for a Molecular Dynamic Simulation of 150 ns.

### Root Mean Square Fluctuation

Root Mean Square Fluctuation displays the flexibility at the residual level (Sangeet et al., 2021). The provided RMS fluctuation (RMSF) graph (Figure 9) illustrates the root mean square fluctuations of the p53 protein, simulated for 150 nanoseconds (ns) under different conditions. The black curve represents the p53 protein without any ligand, showing natural fluctuations with low RMSF values and a few peaks around residues 112 to 125 and 175 to 190, indicating regions interspersed with more flexible segments. The red curve corresponds to the presence of the commercial drug Prima, which induces similar fluctuations as that of clean protein, especially around residue 112 to 125, suggesting increased flexibility in this region and affecting the protein’s overall stability. Prima reduces the fluctuation in the region 175 to 190 compared with the clean protein. Ligand Z3, represented by the green curve, shows a sharp drop around residue 112 to 125, indicating a localized and restricted effect on protein dynamics, while the rest of the curve follows a pattern similar to the Prima, implying Z3’s effect is primarily localized. In contrast, ligand Z7, depicted by the blue curve, shows a higher RMSF values around residue 112 to 125 as compared to Z3 but less than Protein, Prima and Z33. Moreover, Z3 and Z7 induces a lower fluctuation in the active region of the mutant protein (residues 220 to 230) suggesting a restricted dynamic of the active site. The magenta curve for ligand Z33 shows high resembling behaviour with Prima suggesting a similar residual fluctuation effect with moderate flexibility. Overall, the analysis reveals that different ligands induce varying degrees of flexibility and dynamic changes in the p53 protein. Especially Z3 and Z7 induces a restricted motion of active site residues, highlighting their potential for further investigation as therapeutic agents.

**Figure 9:**
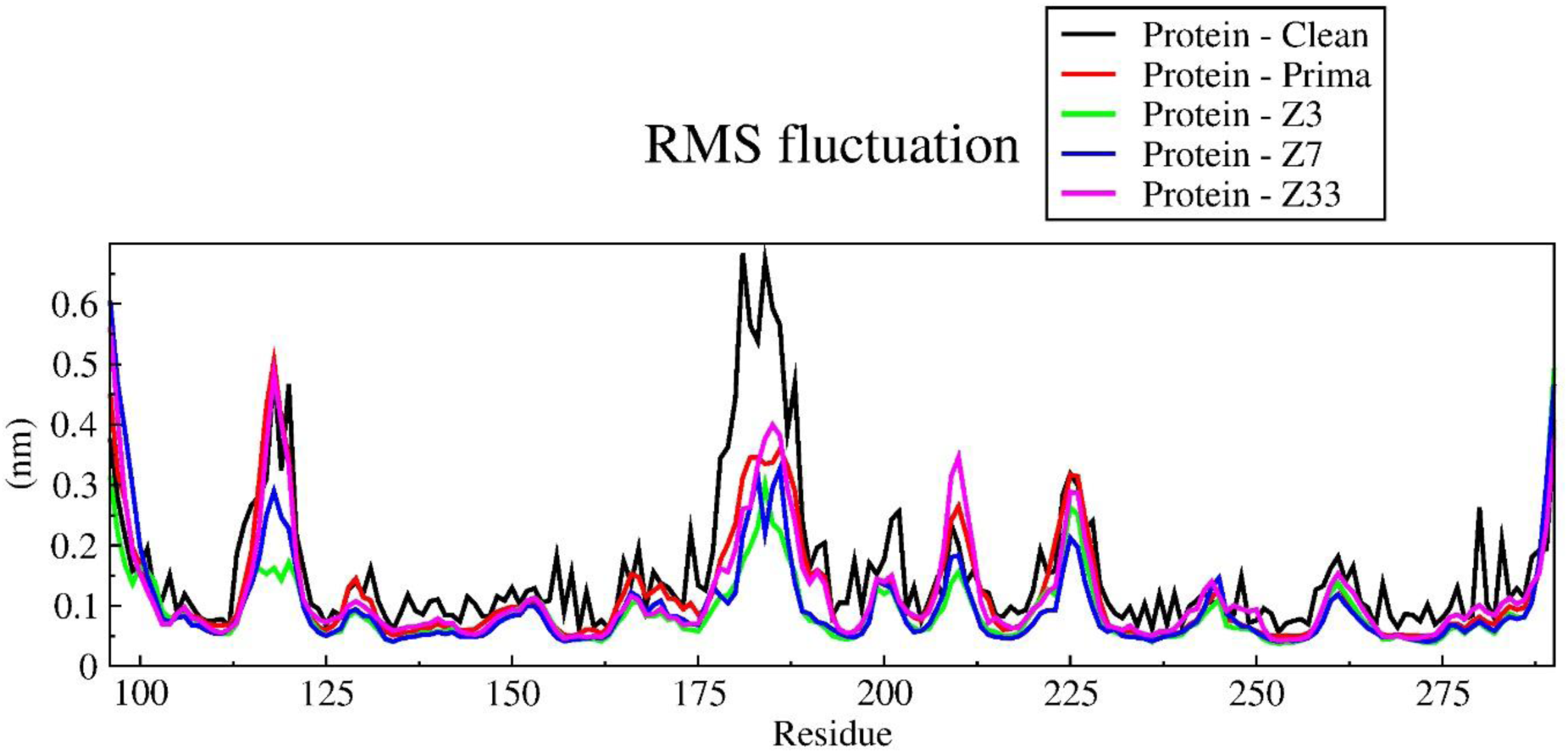
Root Mean Square Fluctuation of Protein without any ligand (Black), Prima (red) and top hit phytochemicals: Z3 (Green), Z7 (Blue) and Z33 (Magenta) for a Molecular Dynamic Simulation of 150 ns.

### MM-PBSA Analysis

The MM-PBSA and MM-GBSA analyses of the ligands binding to the mutant p53 receptor provide valuable insights into the binding energetics. For all the top hit ligands and Prima both methods indicate that while the gas-phase interactions, primarily driven by residues Glu221 (for Prima and Z7), Asn200 and Glu221 (for Z3 and Z33) (Figure 10), favour binding, the solvation energy terms oppose the binding process.

**Figure 10:**
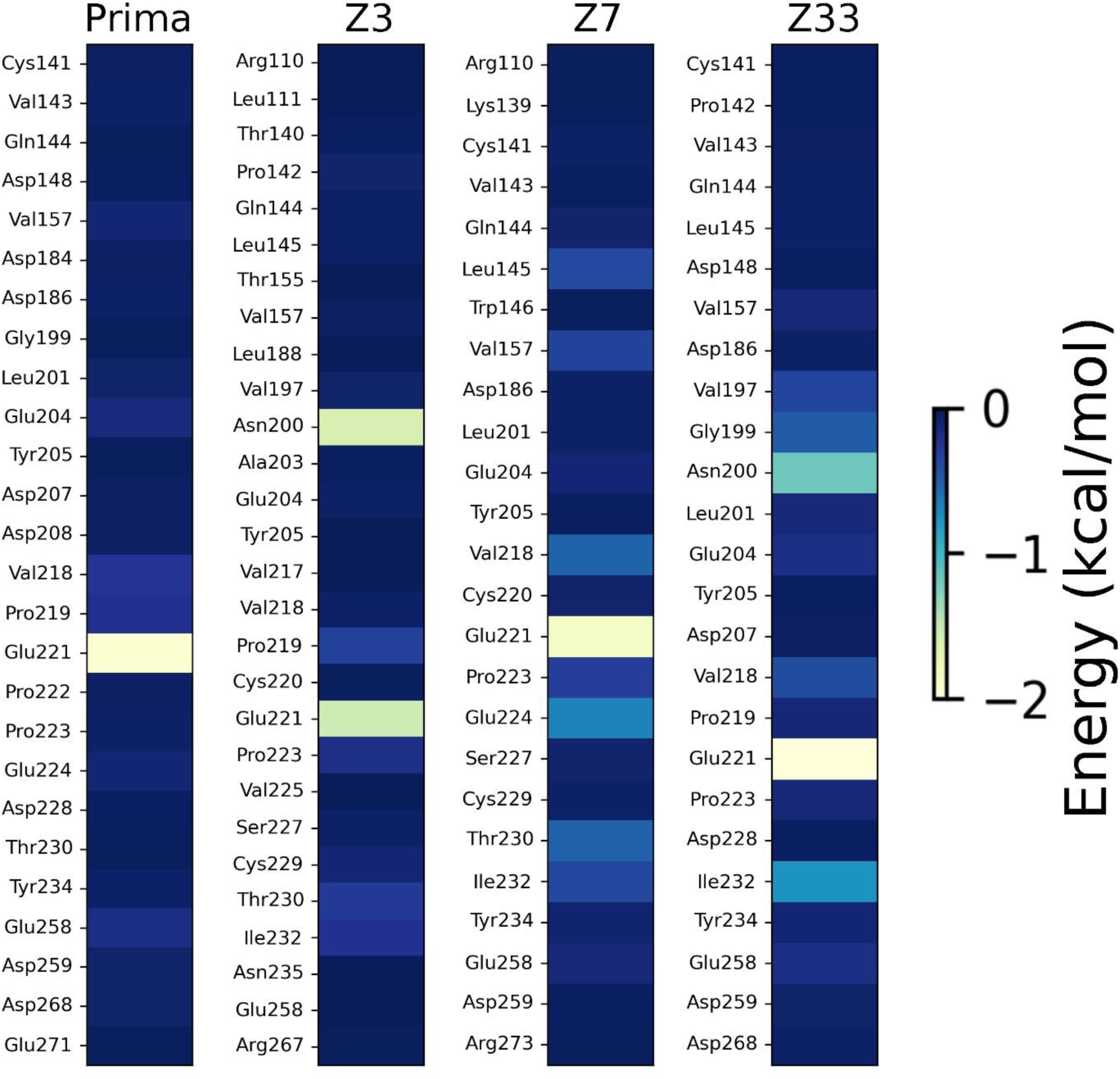
Per residue decomposition energy for Prima and the top hit phytochemicals.

Ultimately, the overall binding energy calculated from these methods is negative and dominated by PB4 (Figure 11) and GB1 (see Supplementary Figure S1) elements, implying a favourable binding affinity for the top hit ligands with the p53 receptor. These computational results suggest that further optimization of these, may be necessary to enhance its binding affinity and potential therapeutic efficacy as a p53 inhibitor.

**Figure 11:**
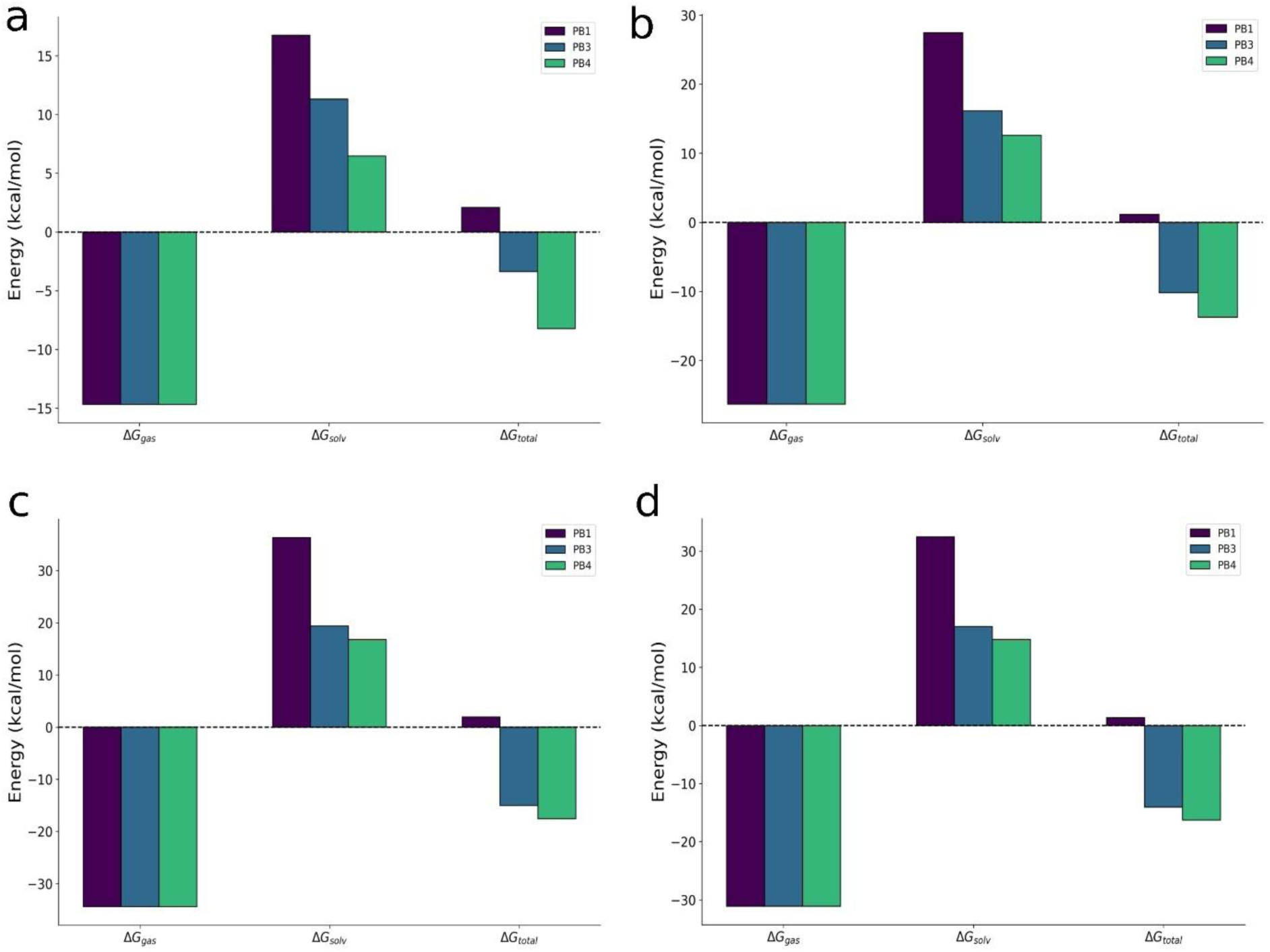
MMPBSA profile of mutant p53 in association with (a) Prima (b) Z3 (c) Z7 and (d) Z33.

## Conclusion

Our comprehensive GC-MS analysis and in *silico* investigation of *Zanthoxylum armatum* DC phytochemicals showed good binding efficacy against the p53 mutant protein involved in throat cancer. The study identified three lead compounds viz. Z3, Z7, and Z33, which demonstrated superior binding affinities and more favourable interaction profiles compared to the commercial drug Prima. The molecular dynamics simulations revealed that these phytochemicals induced more compact and stable protein conformations, with Z3 and Z7 particularly restricting the motion of active site residues. This suggests a potential mechanism for inhibiting the oncogenic functions of mutant p53. The MMPBSA analysis further corroborated these findings, showing favourable binding energetics for the top hit compounds. Importantly, the ADMET analysis indicated that these phytochemicals possess drug-like properties and potentially lower toxicity profiles compared to Prima. This is particularly significant given the challenges associated with off-target effects and toxicity in current synthetic approaches targeting mutant p53. The unique interaction patterns observed, particularly for Z3 and Z7, warrant further investigation into their potential synergistic effects and ability to modulate specific mutant p53 functions. These findings not only contribute to our understanding of mutant p53 targeting but also open new avenues for natural product-based drug discovery in cancer therapeutics. While these results are promising, it is crucial to note that experimental validation is necessary to confirm the efficacy and safety of these compounds. Future studies should focus on in vitro and in vivo experiments to validate the predicted interactions and assess the physiological impacts of these phytochemicals on mutant p53-driven cancer processes. In conclusion, this study provides a strong foundation for the development of novel, nature-inspired therapeutics targeting mutant p53 in throat cancer. The identified lead compounds offer potential alternatives to current synthetic approaches, with the added benefits of potentially reduced toxicity and multi-target effects characteristic of natural products. These findings represent a significant step forward in the quest for more effective and safer treatments for throat cancer and potentially other p53-mutant cancers.

## References

[1] Lorenzoni, V.; Chaturvedi, A.K.; Vignat, J.; Laversanne, M.; Bray, F.; Vaccarella, S. The Current Burden of Oropharyngeal Cancer: A Global Assessment Based on GLOBOCAN 2020. Cancer Epidemiol. Biomarkers Prev., 2022, 31(11):2054–2062. 10.1158/1055-9965.EPI-22-0642

[2] Patterson, R.H.; Fischman, V.G.; Wasserman, I.; Siu, J.; Shrime, M.G.; Fagan, J.F.; Koch, W.; Alkire, B.C. Global Burden of Head and Neck Cancer: Economic Consequences, Health, and the Role of Surgery. Otolaryngol Head Neck Surg., 2020, 162(3):296–303. 10.1177/0194599819897265

[3] Feroz, W.; Sheikh, A.M.A. Exploring the multiple roles of guardian of the genome: P53. Egypt J Med Hum Genet., 2020, 21(49). 10.1186/s43042-020-00089-x

[4] Marei, H.E.; Althani, A.; Afifi, N.; Hasan, A.; Caceci, T.; Pozzoli, G.; Morrione, A.; Giordano, A.; Cenciarelli, C. p53 signalling in cancer progression and therapy. Cancer Cell Int., 2021, 21(1):703. 10.1186/s12935-021-02396-8

[5] Berke, T.P.; Slight, S.H.; Hyder, S.M. Role of Reactivating Mutant p53 Protein in Suppressing Growth and Metastasis of Triple-Negative Breast Cancer. Onco Targets Ther., 2022, 15:23–30. 10.2147/OTT.S342292

[6] Sammons, M.A.; Nguyen, T.T.; McDade, S.S.; Fischer, M. Tumor suppressor p53: from engaging DNA to target gene regulation. Nucleic Acids Res., 2020, 48(16):8848–8869. 10.1093/nar/gkaa666

[7] Nathan, C.A.; Khandelwal, A.R.; Wolf, G.T.; Rodrigo, J.P.; Makitie, A.A.; Saba, N.F.; Forastiere, A.A.; Bradford, C.R.; Ferlito, A. TP53 Mutations in head and neck cancer. Mol Carcinog., 2022, 61(4):385–391. 10.1002/mc.23385

[8] Chen, X.; Zhang, T.; Su, W.; Dou, Z.; Zhao, D.; Jin, X.; Lei, H.; Wang, J.; Xie, X.; Cheng, B.; Li, Q.; Zhang, H.; D, C. Mutant p53 in cancer: from molecular mechanism to therapeutic modulation. Cell Death Dis., 2022, 13(11):974. 10.1038/s41419-022-05408-1

[9] Weisz, L.; Damalas, A.; Liontos, M.; Karakaidos, P.; Fontemaggi, G.; Maor-Aloni, R.; Kalis, M.; Levrero, M.; Strano, S.; Gorgoulis, V.G.; Rotter, V.; Blandino, G.; Oren, M. Mutant p53 Enhances Nuclear Factor kB Activation by Tumor Necrosis Factor α in Cancer Cells. Cancer Res., 2007 67(6):2396–2401. 10.1158/0008-5472.CAN-06-2425

[10] Kikuchi, S.; Nishimura, R.; Osako, T.; Okumura, Y.; Nishiyama, Y.; Toyozumi, Y.; Arima, N. Definition of p53 overexpression and its association with the clinopathological features in luminal/HER2-ngetaive breast cancer. Anticancer Research, 2013, 33(9):3891–3897.

[11] Parrales, A.; Iwakuma, T. Targeting Oncogenic Mutant p53 for Cancer Therapy. Front Oncol, 2015, 5:288. 10.3389/fonc.2015.00288

[12] Duffy, M.J.; Tang, M.; Rajaram, S.; O’Grady, S.; Crown, J. Targeting Mutant p53 for Cancer Treatment: Moving Closer to Clinical Use? Cancers, 2022, 14(18):4499. 10.3390/camcers/14184499

[13] Roszkowska, K. A.; Piecuch, A.; Sady, M.; Gajewski, Z.; Flis, S. Gain of Function (GOF) Mutant p53 in Cancer – Current Therapeutic Approaches. Int J Mol Sci, 2022, 23(21):13287. 10.3390/ijms/232113287

[14] Duffy, M.J.; Synnott, N.C.; McGowan, P.M.; Crown, J.; O’Connor, D.; Gallagher, W.M. p53 as a target for the treatment of cancer. Cancer Treat Rev., 2014, 40(10):1153–1160. 10.1016/j.ctrv.2014.10.004

[15] Izetti, P.; Hautefeuille, A.; Abujamra, A.L.; de Farias, C.B.; Giacomazzi, J.; Alemar, B.; Lenz, G.; Roesler, R.; Schwartsmann, G.; Osvaldt, A.B.; Hainaut, P.; Ashton-Prolla, P. PRIMA-1, a mutant p53 reactivator, induces apoptosis and enhances chemotherapeutic cytotoxicity in pancreatic cancer cell lines. Invest New Drugs, 2014, 32:783–794. 10.1007/s10637-014-0090-9

[16] Hientz, K.; Mohr, A.; Bhakta-Guha, D.; Efferth, T. The role of p53 in cancer drug resistance and targeted chemotherapy. Oncotarget., 2017, 8(5):8921–8946. 10.18632/oncotarget.13475

[17] Zhang, S.; Carlsen, L.; Borrero, L.H.; Seyhan, A.A.; Tian, X.; El-Deiry, W.S. Advanced Startegies for Therapeutic Targeting of Wild-Type and Mutant p53 in Cancer. Biomol., 2022, 12(4):548. 10.3390/biom12040548

[18] Mori, A.; Lehmann, S.; O’Kelly, J.; Kumagai, T.; Desmond, J.C.; Pervan, M.; McBride, W.H.; Kizaki, M.; Koeffler, H.P. Capsaicin, a Component of Red Peppers, Inhibits the Growth of Androgen-Independent, p53 Mutant Prostsate Cancer Cells. Cancer Res., 2006, 66(6):3222–3229. 10.1158/0008-5472.CAN-05-0087

[19] Manna, K.; Das, U.; Das, D.; Kesh, S.B.; Khan, A.; Chakraborty, A.; Dey, S. Naringin inhibits gamma radiation-induced oxidative DNA damage and inflammation, by modulating p53 and NF-kB signalling pathways in murine splenocytes. Free Radic Res., 2015, 49(4):422–439. 10.3109/10715762.2015.1016018

[20] Talib, W.H.; Al-Hadid, S.A.; Ali, M.B.W.; Al-Yasari, I.H.; Ali, M.R.A. Role of curcumin in regulating p53 in breast cancer: an overview of the mechansim of action. Breast cancer (Dove Med Press), 2018, 10:207–217. 10.2147/BCTT.S167812

[21] Javed, Z.; Khan, K.; Herrera-Bravo, J.; Naeem, S.; Iqbal, M.J.; Sadia, H.; Qadri, Q.R.; Raza, S.; Irshad, A.; Akbar, A.; Reiner, Ž.; Al-Harrasi, A.; Al-Rawahi, A.; Satmbekova, D.; Butnariu, M.; Bagiu, I.C.; Bagiu, R.V.; Sharifi-Rad, J. Genistein as a regulator of signalling pathways and microRNAs in different types of cancers. Cancer Cell Int, 2021, 22:256. 10.1186/s12935-022-02667-y

[22] Talib, W.H.; Awajan, D.; Alqudah, A.; Alsawwaf, R.; Althunibat, R.; Abu AlRoos, M.; Al Safadi, A.; Abu Asab, S.; Hadi, R.W.; Al Kury, L.T. Targeting Cancer Hallmarks with Epigallocatechin Gallate (EGCG): Mechanistic Basis and Therapeutic Targets. Molecules, 2024, 29(6):1373. 10.3390/molecules29061373

[23] Singh, T.D.; Meitei, H.T.; Sharma, A.L.; Robinson, A.; Singh, L.S.; Singh, T.R. Anticancer propertieis and enhancement of therapeutic potential of cisplatin by leaf extract of Zanthoxylum armatum DC. Biol Res, 2015, 48(1):46. 10.1186/s40659-015-0037-4

[24] Okagu, I.U.; Ndefo, J.C.; Aham, E.C.; Udenigwe, C.C. Zanthoxylum Species: A Review of Traditional Uses, Phytochemistry and Pharmacology in Relation to Cancer, Infectious Diseases and Sickle Cell Anemia. Front Pharmacol, 2021, 12:713090. 10.3389/fphar.2021.713090

[25] Alam, F.; Najum us Saqib, Q.; Waheed, A. Cytotoxic activity of extracts and crude saponins from Zanthoxylum armatum DC. Against human breast (MCF-7, MDA-MB-468) and colorectal (Caco-2) cancer cell lines. BMC Complement Altern Med, 2017, 17(1):368. 10.1186/s12906-017-1882-1

[26] Morris, G.M.; Huey, R.; Lindstrom, W.; Sanner, M.F.; Belew, R.K.; Goodsell, D.S.; Olson, A.J. Autodock4 and AutoDockTools4: automated docking with selective receptor flexibility. J. Comput Chem., 2009, 30(16):2785–2791. 10.1002/jcc.21256

[27] Sangeet, S.; Khan, A. An in-silico approach to identify bioactive phytochemicals from *Houttuynia cordata* Thunb. As potential inhibitors of human glutathione reductase. Journal of Biomolecular Structure and Dynamics, 18:1–20. 10.1080/07391102.2023.2294181

[28] Daina, A.; Michielin, O.; Zoete, V. SwissADME: a free web tool to evaluate pharmacokinetics, drug-likeness and medicinal chemistry friendliness of small molecules. Sci Rep., 2017, 7:42717. 10.1038/srep42717

[29] Banerjee, P.; Kemmler, E.; Dunkel, M.; Preissner, R. ProTox 3.0: a webserver for the prediction of toxicity of chemicals. Nucleic Acids Res., 52(W1):W513–W520. 10.1093/nar/gkae303

[30] Abraham, M. J.; Murtola, T.; Schulz, R.; Páll, S.; Smith, J. C.; Hess, B.; Lindahl, E. Gromacs: High performance molecular simulations through multi-level parallelism from laptops to supercomputers. SoftwareX, 2015, 1(2):19–25. 10.1016/j.softx.2015.06.001

[31] Hess, B.; Kutzner, C.; van der Spoel, D.; Lindahl, E. GROMACS 4: Algorithms for highly efficient, load-balanced and scalable molecular simulation. J Chem Comput, 2008, 4(3): 435–447. 10.1021/CT700301Q

[32] Van Der Spoel, D.; Lindahl, E.; Hess, B.; Groenhof, G.; Mark, A. E.; Berendsen, H. J. C. GROMACS: Fast, flexible, and free. J Comput Chem, 2005, 26(16):1701–1718. 10.1002/JCC.20291

[33] Vanommeslaeghe, K.; Hatcher, E.; Acharya, C.; Kundu, S.; Zhong, S.; Shim, J.; Darian, E.; Guvench, O.; Lopes, P.; Vorobyov, I.; MacKerell, A. D. Jr. CHARMM General Force Field (CGenFF): A force field for drug-like molecules compatible with the CHARMM all-atom additive biological force fields. J Comput Chem, 2010, 31(4):671–690. 10.1002/jcc.21367

[34] Sangeet, S.; Khan, A.; Prabha, S.; Pandey, K. M. Antibacterial property of biologically synthesised iron nanoparticles against Staphylococcus aureus. In P. Verma, O. D. Samuel, T. N. Verma, & G. Dwivedi (Eds.), Advancement in materials, manufacturing and energy engineering, Vol. II. Lecture notes in mechanical engineering, Springer, Singapore, 2022. 10.1007/978-981-16-8341-1_7

[35] Wang, Z.; Pan, H.; Sun, H.; Kang, Y.; Liu, H.; Cao, D.; Hou, T. (2022). fastDRH: A webserver to predict and analyse complexes based on molecular docking and MM/PB(GB)SA computation. Breif. Bioinform, 23(5):bbac201. 10.1093/bib/bbac201

[36] Zeenat, L.; Prajapati, S.; Sangeet, S.; Khan, A; Pandey, K.M. In-silico screening of potential phytochemicals against extracellular adherence (EAP) protein of *Staphylococcus aureus* from Indian Medicinal Plants. Research Journal of Pharmacy and Technology, 2023, 16(10):4691–4697. 10.52711/0974-360X.2023.00762

[37] Sangeet, S.; Khan, A.; Mahanta, S.; Roy, N.; Das, S.K.; Mohanta, Y.K.; Saravanan, M.; Tag, H.; Hui, P.K. Computational Analysis of *Bacopa monnieri* (L.) Wettst. Compounds for Drug Discovery against Neurodegenerative Disorders. Curr Comput Aided Drug Des., 2022, 19(1):24–36. 10.2174/1573409918666221010103652

[38] Menichini, P.; Monti, P.; Speciale, A.; Cutrona, G.; Matis, S.; Fais, F.; Taiana, E.; Neri, A.; Bomben, R.; Gentile, M.; Gattei, V.; Ferrarini, M.; Morabito, F.; Fronza, G. Antitumour Effects of PRIMA-1 and PRIMA-1^Met^ (APR246) in Haematological Malignancies: Still a Mutant P53-Dependent Affair? Cells, 2021, 10(1):98. 10.3390/cells10010098

