## Supplemental Table 1 and Figure s1 for "Computational Screening of *Zanthoxylum armatum* DC Phytochemicals for inhibiting Mutant P53 in Throat Cancer Therapy"

**Table 1:** Lipinski's Parameter for the phytochemicals of *Zanthoxylum armatum* DC

| Phytochemical | | Lipinski Parameters | | | | | |
|---|---|---|---|---|---|---|---|
| **Code** | **Compound Name** | **MW** | **LogP** | **HD** | **HA** | **V** | **LD50** |
| Z1 | (1R,2R,10S,11R,13S,15R)-11-hydroxy-15-methyl-6-azatetracyclo[8.6.0.01,6.02,13]hexadecan-14-one | 263.38 | 1.8396 | 1 | 3 | No | 200 |
| Z2 | (3R,5S,7R,8R,9S,10S,12S,13R,14S,17R)-17-[(2R)-5,6-dihydroxyheptan-2-yl]-10,13-dimethyl-2,3,4,5,6,7,8,9,11,12,14,15,16,17-tetradecahydro-1H-cyclopenta[a]phenanthrene-3,7,12-triol | 438.64 | 3.1058 | 5 | 5 | No | 500 |
| Z3 | (8R,9S,13S,14S,17S)-13,17-dimethyl-7,8,9,11,12,14,15,16-octahydro-6H-cyclopenta[a]phenanthrene-3,17-diol | 286.41 | 3.9993 | 2 | 2 | No | 2000 |
| Z4 | 2-amino-10H-acridin-9-one | 210.23 | 2.2634 | 2 | 2 | No | 360 |
| Z5 | (1R,9S,15S)-15-amino-1-methyltricyclo[7.5.1.02,7]pentadeca-2(7),3,5-trien-4-ol | 245.36 | 3.1136 | 2 | 2 | No | 243 |
| Z6 | (2E,4E,8E,10E,12E)-N-(2-methylpropyl)tetradeca-2,4,8,10,12-pentaenamide | 273.42 | 4.3397 | 1 | 1 | No | 2000 |
| Z7 | (3R,5R,7R,8R,9S,10S,12S,13R,14S,17R)-17-[(2R)-5-hydroxypentan-2-yl]-10,13-dimethyl-2,3,4,5,6,7,8,9,11,12,14,15,16,17-tetradecahydro-1H-cyclopenta[a]phenanthrene-3,7,12-triol | 394.59 | 3.3564 | 4 | 4 | No | 2000 |
| Z8 | (E)-3-(1H-indol-3-yl)prop-2-enoic acid | 187.19 | 2.2657 | 2 | 1 | No | 2500 |
| Z9 | (E,3R,5S)-7-[3-(4-fluorophenyl)-1-propan-2-ylindol-2-yl]-3,5-dihydroxyhept-6-enoic acid | 411.473 | 4.6281 | 3 | 4 | No | 416 |
| Z10 | 6,7-dimethoxy-1-[(4-methoxyphenyl)methyl]-1,2,3,4-tetrahydroisoquinolin-5-ol | 329.396 | 2.8475 | 2 | 5 | No | 2000 |
| Z11 | (8S)-8-(2-hydroxypropan-2-yl)-8,9-dihydrofuro[2,3-h]chromen-2-one | 246.262 | 1.8674 | 1 | 4 | No | 832 |
| Z12 | (1R,2R,8R,10R,13S,15R)-8-hydroxy-15-methyl-6-azapentacyclo[8.6.0.01,6.02,13.04,10]hexadecan-11-one | 261.364 | 1.4467 | 1 | 3 | No | 400 |
| Z13 | 7-hydroxy-3-[4-hydroxy-3-(3-methylbut-2-enyl)phenyl]chromen-4-one | 322.36 | 4.3799 | 2 | 4 | No | 2500 |
| Z14 | (3R,5R,8R,9R,10S,13S,14S)-3-hydroxy-13-methyl-2,3,4,5,6,7,8,9,10,11,12,14,15,16-tetradecahydro-1H-cyclopenta[a]phenanthren-17-one | 276.419 | 3.5690 | 1 | 2 | No | 3000 |
| Z15 | 4-(4-hydroxyphenyl)phenol | 186.209 | 2.7648 | 2 | 2 | No | 4920 |
| Z16 | (2S,3S,6S)-6-[(1E,3E,5E,7E)-deca-1,3,5,7-tetraenyl]-1,2-dimethylpiperidin-3-ol | 261.408 9 | 3.4648 | 1 | 2 | No | 3500 |
| Z17 | 1-methyl-4,9-dihydro-3H-pyrido[3,4-b]indol-7-ol | 200.241 | 2.2386 | 2 | 2 | No | 445 |
| Z18 | Ethyl 2-acetamido-3-(4-hydroxyphenyl)propanoate | 251.282 | 1.0025 | 2 | 4 | No | 7710 |
| Z19 | 2-heptyl-3-hydroxy-1H-quinolin-4-one | 259.349 | 3.7466 | 2 | 2 | No | 753 |
| Z20 | Carvone oxide | 166.219 | 1.6991 | 0 | 2 | No | 5000 |
| Z21 | 8,8a-Deoxyoleandolide | 372.502 | 1.7901 | 3 | 6 | No | 5000 |
| Z22 | Homodolicholide | 492.697 | 3.7002 | 4 | 6 | No | 1000 |
| Z23 | 17alpha-Methylestradiol | 286.414 | 3.9993 | 2 | 2 | No | 2000 |
| Z24 | Dezocine | 245.365 | 3.1136 | 2 | 2 | No | 243 |
| Z25 | Colchicine | 399.443 | 2.8716 | 1 | 6 | No | 6 |
| Z26 | xi-Anomuricine | 329.396 | 2.8475 | 2 | 5 | No | 2000 |

| | | | | | | | |
|---|---|---|---|---|---|---|---|
| Z27 | Thiofanox | 218.321 | 2.1075 | 1 | 4 | No | 153 |
| Z28 | Fawcettidine | 245.365 | 2.9913 | 0 | 2 | No | 1273 |
| Z29 | 27-Nor-5b-cholestane3a,7a,12a,24,25-pentol | 438.649 | 3.1058 | 5 | 5 | No | 500 |
| Z30 | Harmalol | 200.241 | 2.2386 | 2 | 2 | No | 445 |
| Z31 | Armillarin | 414.498 | 3.5670 | 2 | 6 | No | 500 |
| Z32 | 2-Heptyl-3-hydroxy-quinolone | 259.349 | 3.7466 | 2 | 2 | No | 753 |
| Z33 | Annotinine | 275.347 | 1.1897 | 0 | 4 | No | 3000 |
| Z34 | Ruspolinone | 249.309 | 2.0286 | 1 | 4 | No | 656 |
| Z35 | alpha-Eucaine | 333.248 | 3.0380 | 0 | 5 | No | 2 |
| Z36 | Cryptophorine | 261.408 | 3.4648 | 1 | 2 | No | 3500 |
| Z37 | 2,2-Dimethyl-3-(4- methoxyphenyl)-4-propyl-2H1-benzopyran-7-ol | 366.457 | 5.5024 | 0 | 4 | Yes | 500 |
| Z38 | Cuauchichicine | 343.511 | 3.8664 | 0 | 3 | No | 500 |
| Z39 | 2-Hydroxy-6-oxo-octa-2,4- dienoate | 418.486 | 5.0121 | 2 | 3 | Yes | 1650 |
| Z40 | Quinic acid | 192.167 | -2.3213 | 5 | 5 | No | 9800 |
| Z41 | Crotanecine | 171.195 | -1.6753 | 3 | 4 | No | 3500 |
| Z42 | Heliotrine | 313.394 0 | 0.32679 | 2 | 6 | No | 46 |
| Z43 | 3-Methoxymorphinan | 257.376 | 3.0412 | 1 | 2 | No | 116 |

*MW – Molecular Weight*

*HD – Number of Hydrogen Donor*

*HA – Number of Hydrogen Acceptor*

*V – Violation*

*LD50 – in units of mg/kg*

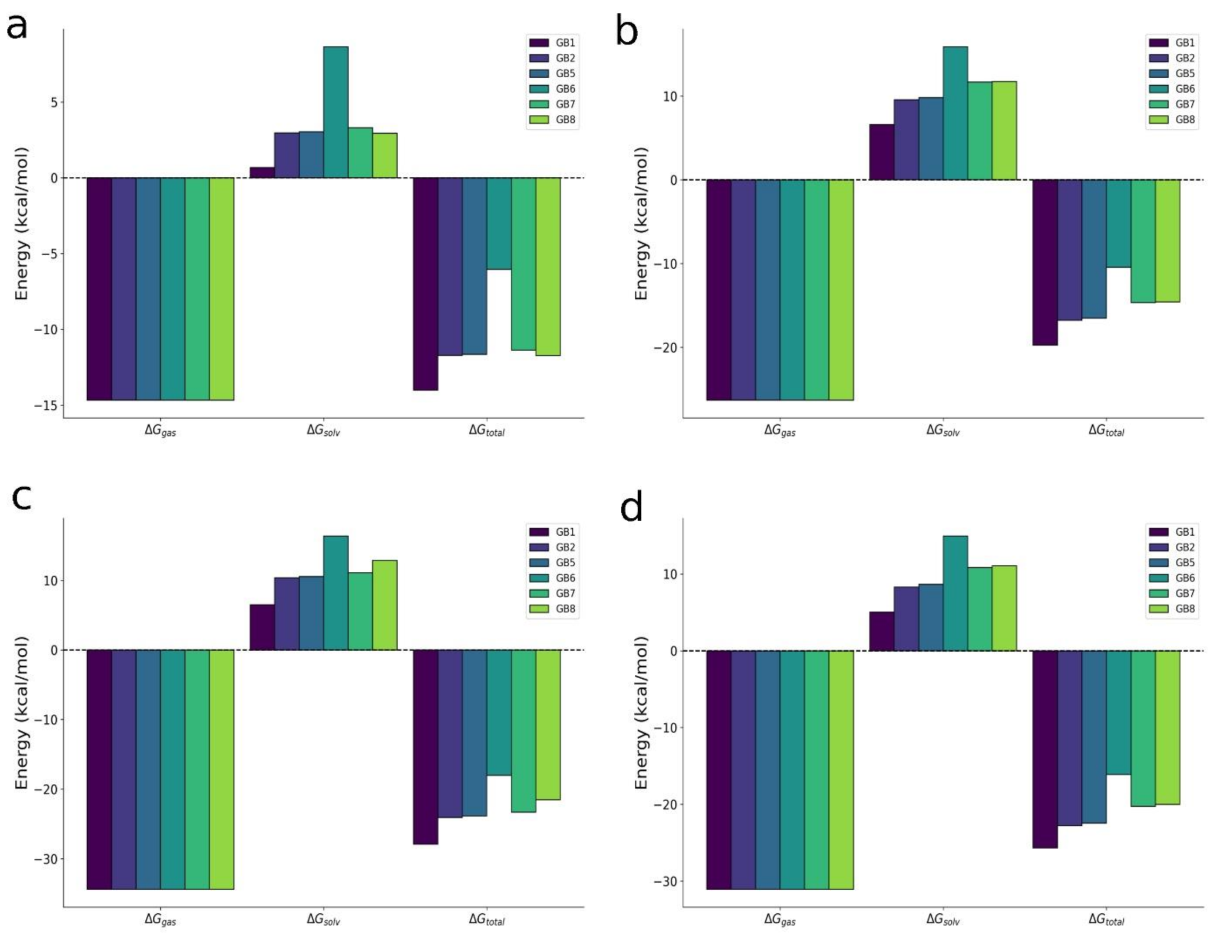


**Figure S1**: MMGBSA profile of mutant p53 in association with (a) Prima (b) Z3 (c) Z7 and (d) Z33
